# Porcine extended pluripotent stem cell-derived somite-like mesoderm cells with Dox-driven PAX7 are robust seed cell resource for facilitating production of cultured meat

**DOI:** 10.64898/2026.03.03.709441

**Authors:** Cong Xia, Sheng Ye, Hai Wang, Meichao Wang, Renquan Zhang, Hongliang Yu, Lijia Wen, Xueliang Wang, Yinghua Ye, Xiner Feng, Bingxiu Ma, Han Wu, Liangxue Lai

**Affiliations:** CAS Key Laboratory of Regenerative Biology, Guangdong Provincial Key Laboratory of Stem Cell and Regenerative Medicine, Guangzhou Institutes of Biomedicine and Health, Chinese Academy of Sciences, Guangzhou 510530, China; University of Chinese Academy of Sciences, Beijing 100049, China; Sanya Institute of Swine Resource, Hainan Provincial Research Centre of Laboratory Animals, Sanya 572000, China; Guangdong Provincial Key Laboratory of Large Animal Models for Biomedicine, Wuyi University, Jiangmen 529020, China; Key Laboratory of Zoonosis Research, Ministry of Education, College of Animal Science, Jilin University, Changchun 130062, China; Department of Hepatobiliary and Pancreatic Surgery, General Surgery Center, The First Hospital of Jilin University, Changchun 130021, China

**Keywords:** Somite-liKe mesoderm cells, Doxycycline-inducible PAX7, Long-term passage, 3D scaffold, Cultured meat

## Abstract

Cultured meat technology, with its significant advantages of shortening meat production cycles, reducing natural resource consumption, minimizing the risk of zoonotic disease transmission, and enabling precise control over nutritional composition and texture, offers a novel alternative source for human meat consumption. One of the major challenges to produce cultured meat in large scale is how to establish high.quality seed cells, which should have long term proliferative capacities and are able to differentiate into muscles efficienuy with simple procedures. Here, we first established an engineered porcine expanded potential stem cells (Tet-On-PAX7 EPSCs) containing Tet-On regulated PAX7 gene. Then the Tet-On-PAX7 EPSCs were induced to somite-liKe mesodermal cells. These somite-liKe mesodermal cells can be expanded over 10^25^-fold even after 40 passages *in-vitro* culture while retaining strong myogenic potential. The somite-like mesodermal cells treated with DOX for one day would differentiate into muscle stem cells (Muses), and the later were able to differentiate into muscles with an efficiency of up to 90% within just 7 days in 11-FSDeDa without Dox. Moreover, when somite-liKe mesodermal cells were seeded on patterned scaffolds, microcarrier scaffolds, or cultured in anchorage-independent suspension, they maintained high efficiency in muscle differentiation, confirming their potential to be used as seed cells for scaled cultured meat production.

## 1. Introduction

With the global population growing, the demand for meat products continues to expand (Stoll-Kleemann and O’Riordan, 2015). However, traditional livestocK farming not only exerts pressure on the environment (Tuomisto and de Mattos, 2011), but contributes to the transmission of zoonotic diseases (Hsi et al., 2015; Wang et al., 2018) and the overuse of antibiotics (McCracKin et al., 2016). The cultured meat technology mimics the development and injury repair mechanism of animal muscles *in-vitro,* bypassing the traditional breeding process and directly producing meat products through scalable bioprocessing of cells (Stephens et al., 2018), offering a new pathway for efficient meat supply with shortened production cycles. Furthermore, the flavor and nutritional value of cultured meat can be adjusted through precise regulation of its components (Elhaddad et al., 2025). Therefore, this technology hOlds significant potential to meet the growing global demand for clean meat {Bhat et al., 2017). The rapid development of the cultured meat industry is marKed by regulatory approvals granted to the products from multiple companies in countries such as the United States, Singapore, and Israel (Choi et al., 2025).

For production of cultured meat in large-scale, one of the major challenges is how to establish high-quality seed cells, which should have long term proliferative capacities and are able to differentiate into muscles efficiently with simple procedures (Xie et al., 2025). So far, the three main types of seed cells including adult stem cells, immortalized cell lines, and embryo-derived pluripotent stem cells, have been used for the study of cultured meat (Martins et al., 2024). Muscle satellite cells, also referred to as muscle stem cells (MuSC), are often regarded as the primary cell source due to their high muscle-forming potential (Kadim et al., 2015; Li et al., 2015). However, these cells reside between the sarcolemma and basement membrane (Forcina et al., 2019; Musaro and Carosio, 2017), thusacquisition of these cells still relies on animal tissues(Krishnan et al., 2026; Wang et al., 2025; Yun et al., 2023), which runs counter to the core philosophy of cultured meat. In addition, Muses exhibit the HayflicK limit-a finite proliferative capacity-and gradually lose their differentiation potential (Dan-Jumbo et al., 2024; Messmer et al., 2022; Redondo et al., 2017). Moreover, these cells require isolation through antibody-based sorting methods (Motohashi et al., 2014), a complex procedure with high costs. Immortalized cell lines established through spontaneous immortalization (PasitKa et al., 2023) or genetic engineering (FlacK, 2025), possessing sustained proliferative capacity, have been used as the seed cells for cultured meat. However inherent metabolic characteristics and protein expression profiles of such cells differ from those of genuine myocytes, and would result in non-muscle-directional differentiation, often impacting the accurate flavor and texture of cultured meat product.

Pluripotent stem cells including embryonic stem cells (ESCs) and induced pluripotent stem cells (iPSCs) are considered as highly promising seed cells for production of cultured meat (Kumar et al., 2021) since they not only possess the capacities of unlimited proliferation and differentiation into diverse cell lineages, but also maintain the genomic stability. This concept had been proved in porcine cultured meat by Han’s group, in which porcine pre-gastrulation epiblast stem cells (pgEpiSCs) were used to develop a directional differentiation system for generating sKeletal muscle fibers with typical muscular characteristics and transcriptional features during myogenic differentiation (Zhu et al., 2023). However, the myogenic differentiation process of pgEpiSCs is more complex than that of Muses or immortalized myoblast cell lines (such as bovine satellite cells, chicKen fibroblasts or C2C12) and taKes more than 37 days to achieve terminal differentiation into of myotube, which needs to be simplified or standardized to meet the requirements for fast production process of cultured meat.

In this study, we integrated a Tet-On inducible system (Jin et al., 2023) capable of expressing porcine Paired box 7(PAX7) gene into the pEPSC genome (Tet-On-PAX7 EPSCs), attempting to simplify the process of porcine pluripotent cell differentiating into myotube. We first induced the Tet-On-PAX7 EPSC into somite-liKe mesoderm cells, which can be expanded over 10^25^- fold under prolonged *in-vitro* culture while retaining strong myogenic potential. Using these somite-liKe mesoderm cells as “seed cells”, upon doxycycline (Dox) treatment for one day to drive PAX7 expression, a high efficiency of myotube formation was yielded in just 7 days. In addition to 20 culture, the cells successfully underwent differentiation under multiple 30 differentiation system for shaping cultured tissue, including patterned scaffolds, microcarrier-based systems, and anchorage-independent suspension system. This versatile and scalable precursor seed cell platform provides a compelling alternative to traditional myogenic cell sources and offers promising potential for application in large-scale cultured meat production.

## 2. Materials and methods

### 2.1 Genetic modification of porcine expanded potential stem cells

pEPSCs were seeded onto mitomycin C {MedChemExpress Cat.NO: HY-13316)-inactivated mouse embryonic fibroblast (STO) feeder layers. When the cells reached 70-90% confluency, 10% fetal bovine serum (FBS, Bio-Channel Cat.NO: BC-SE-FBS01) was added to the EPSC medium. After 4 hours, electrotransfection was performed using the Neon® Transfection System (Thermo Fisher Scientific) under the following parameters: 1200 V, 20 ms, 1 pulse. Within 12-24 hours, the EPSC medium containing Y-27632 and 10% FBS was replaced with standard EPSC medium. After 5-7 days of culture, Id-Tomato-positive pEPSC colonies were picKed under a fluorescence microscope using a 1O µL pipette and transferred to a 48-well plate. The following day, the cells were dissociated and reseeded into a 6 cm dish. Once the pEPSC colonies had expanded sufficienuy, pure positive clones were isolated under fluorescence microscopy again using a 1O µL pipette.

### 2.2 Alkaline phosphatase (AP) Staining

After fixation with paraformaldehyde (PFA, Leagene Cat. NO: DF0135), cells were stained following the instructions of the BCIP/NBT AIKaline PhOsphatase Color Development Kit (Beyotime Cat. NO: C3206). The worKing solution was added and incubated at room temperature in the darK for 30 minutes. The staining solution was then removed, and the cells were washed 1-2 times with distilled water. Finally, the results were observed under microscope.

### 2.3 Comet assay

The layer of gel was prepared by coating a glass slide with 1% agarose solution (GenStar Cat. NO: VA10252). Subsequently, cells were digested with trypsin and mixed with a certain volume of 0.75% low-density agarose (Aladdin Cat. NO: A104063) solution to form a second layer, which was overlaid onto the first gel layer. A third gel layer was then applied using 1% agarose solution. Following this, the sampie was lysed at 4°c using a lysis buffer composed of 90% Lysis Buffer (Beyotime Cat. NO: C2041S) and 10% Dimethyl sulfoxide (DMSO Sigma Cat. NO: 04540). DNA was then denatured, subjected to electrophoresis, neutralized, and finally stained with SYRB Green (Solarbio Cat. NO: G8140-1ml). For the positive control, cells were treated with 3% H,O_2_ for 20 minutes.

### 2.4 Reverse transcription quantitative polymerase chain reaction

Total RNA was extracted from cells using the RNA Extraction Kit (Magen, Cat. NO: R4012-03). Reverse transcription of the extracted RNA into cONA was performed using the HiScript **Ill** RT SuperMix for qPCR (+gONA wiper) (Vazyme Cat. NO: R323). The resulting cONA was diluted 10-fold and subjected to quantitative real-time PCR using Chama Universal SYBR qPCR Master Mix (Vazyme Cat. NO: Q711-03). The primers used are listed in Table S1.

### 2.5 Karyotype analysis

Cells were seeded into 6-cm dishes and cultured until they reached approximately 80% confluency. Colchicine (Med Chem Express Cat. NO: HY-16569) was added to the culture medium and incubated for 3 hours. Subsequently, the cells were digested, centrifuged, and the supernatant was discarded. After hypotonic treatment and fixation with PFA, the cells were mounted on slides and stained with Giemsa solution. The slides were gently rinsed with PBS on both sides and air-dried at room temperature. Once dried, metaphase spreads were identified under a microscope. Well-spread chromosomes were observed under oil immersion lens, captured using a chromosome analysis system, and systematically arranged for Karyotypic evaluation.

### 2.6 lmmunofluorescence staining

After fixation with PFA for 1O minutes, the cells were washed with PBS. The samples were treated with a solution containing 5% BSA (Biosharp Cat. NO: A-4612) and 0.3% Triton X-100 (Beyotime Cat. NO: P0096-100ml) for 20 minutes and then washed with PBS. Subsequently, the cells were incubated with primary antibodies overnight at 4°c. After washing, corresponding fluorescent secondary antibodies were applied and incubated for 2 hours at room temperature. Finally, the nuclei were stained with DAPI. Images were captured using a fluorescence microscope. The antibodies used are listed in Table S2.

### 2.7 Flow cytometry

Cells were prepared as single-cell suspensions at-1×106 cells/ml in PBS with 1% BSA. For surface staining, cells were incubated with fluorochrome-conjugated antibodies (Table S2) for 30 minutes at 4°C in the darK. Unbound antibodies were removed by washing with PBS/1% BSA. Cells were resuspended in PBS/1% BSA, filtered through a cell strainer, and acquired on a flow cytometer.

### 2.8 Derivation of somite-like mesoderm cells and method for myogenic differentiation

Tet-On-PAX7 EPSCs were cultured in EPSC medium until they reached 70-80% confluency, and then switched to ICS-ElnDe medium for at least 12 days. The ICS-ElnDe medium consisted of **IMDM** (Gibco Cat. NO: 31980030), 5% HS (Gibco Cat. NO: 26050088), 1% penicillin-streptomycin (Thermo Cat. NO: 15140-122), 3 **µM** CHIR99021 (SellecK Cat. NO: S1263), 2 **µM** SB431542 (APExBIO Cat. NO: A8249), 10 ng/ml hr-EGF (PeproTech Cat. NO: AF-100-15), 10 µg/ml Insulin (SellecK Cat. NO: S6955), 10 **µM** Dexamethasone (Sigma Cat. NO: 04902), 50 µg/ml L-ascorbicacid (Sigma Cat. NO: A4544), and 1 **µM** Y-27632 (SellecK Cat. NO: S1049). This stage represents the differentiation of pEPSCs into somite-liKe mesoderm cells. During this period, when cells reached full confluency, they were passaged onto gelatin-coated dishes without feeder layers. Subsequently, the cells were cultured for 1 day in 11-FSDeDa medium supplemented with 1 µg/ml Doxycycline hyclate (Sigma Cat. NO: 09891). The **11-**FSDeDa medium contained **IMDM** (Gibco Cat. NO: 31980030), 15% KSR (Thermo Cat. NO: 10828028), 10 ng/ml IGF-1 (PeproTech Cat. NO: 250-19), 10 **µM** ForsKolin (SellecK Cat.NO: S2449), 10 **µM** SB431542 (APExBIO Cat. NO: A8249), 10 **µM** Dexamethasone (Sigma Cat.NO: 04902), and 1 µM DAPT (APExBIO Cat. NO: A8200). Finally, Dox was withdrawn, and the cells were further cultured for 7 days, during which spontaneous and irregular myotube contractions could be observed.

### 2.9 Myogenic differentiation procedure of somite-like mesoderm on patterned scaffolds, microcarrier scaffolds, and in anchorage-independent suspension culture

For patterned scaffold differentiation, molds were fabricated via 30 printing, infused with 4% gelatin solution, and cryopreserved at-2o•c overnight. Cells pre-cultured in ICS-ElnDe medium were induced with Dox-containing 11-FSDeDa medium for 1 day, then digested and seeded onto gelatin-patterned scaffolds at 6x 1Cl6 cells/16 cm•. Differentiation proceeded for 7 days post-Dox withdrawal.

In microcarrier scaffold differentiation, 20 mg of Seplife LX-MC-Dex1 microcarriers (100-180 µm diameter) were hydrated in PBS for ≥6 hOurs. Cells were seeded at 2x 1Cl6 cells/20 mg carriers in a 250 ml spinner flasK under dynamic agitation (30 rpm/7 min → O rpm/120 min, 24 cycles → 40 rpm continuous). After 24 hours, complete cell attachment was confirmed by the absence of free-floating cells in the ICS-ElnDe culture medium. Attached cells were then treated with Dox-containing 11-FSDeDa medium for 1 days, followed by 7-day differentiation after Dox removal.

For anchorage-independent suspension differentiation, cells were seeded at 5×10^5^ cells/well in uncoated 6-well plates under orbital shaKing (11O rpm). Cell aggregates (≈50 µm diameter) formed on Day 1, coalesced into larger clusters (≈200 µm) by Days 2-3, and stabilized into compact structures by Days 6-7. Aggregates were then induced with Dox-containing 11-FSDeDa medium for 1 days and differentiated for 7 days after Dox withdrawal.

### 2.10 RNA-seq Analysis

#### 2.10.1 Library preparation and sequencing

RNA extraction, library construction, and high-throughput sequencing were performed by TsingKe Biotechnology Co., ltd. The quality of the extracted RNA was assessed using a NanoOrop spectrophotometer and an Agilent 2100 Bioanalyzer to ensure it met the requirements for library construction. All RNA sampies satisfied the following criteria: total amount ≥ 2 µg (sufficient for two library constructions), concentration ≥ 40 ng/µL, volume ≥ 1O µL, 00260/280 ratio between 1.7 and 2.5, 00260/230 ratio between 0.5 and 2.5, and a normal absorption peaK at 260 nm. Following quality control, libraries were constructed using the lllumina TruSeq RNA Library Preparation Kit. After passing library quality control, paired-end 150 bp (PE150) sequencing was performed on an lllumina NovaSeq X Plus platform. A total of 12 sampies were sequenced in this study. These sampies were divided into four experimental conditions: Tet-On-PAX7-EPSC, EPSC-SMC, EPSC-MuSC, and EPSC-MC, with three biological replicates per condition.

#### 2.10.2 Raw data processing andtranscriptome assembly

The raw sequencing data in FASTQ format were initially processed using Fastp (v0.20.1) for quality control. The resulting high-quality clean reads were then aligned to the reference genome (Sscrofa11.1, GenBanK accession number: GCA_000003025.6) using HISAT2 (v2.2.1). Following alignment, transcriptomes were *de novo* assembled using StringTie (v2.0.4). The assembly results were compared with the reference genome annotation file using Gffcompare (v0.9.8) to obtain the final transcript annotations. Gene expression levels were quantified using FPKM through the ballgown R pacKage, resulting in a gene expression matrix.

#### 2.10.3 Principal component analysis (PCA)

Differential expression analysis was performed on the raw expression matrix using DESeq2 (v1.26.0), which internally applies Relative Log Expression (RLE) normalization for data standardization. Genes with a fold change > 2 (llog_2_Fq > 1) and an adjusted p-value {adj.P) < 0.05 after multiple testing correction were defined as significantly differentially expressed. Subsequently, the transformed expression matrix of these differentially expressed genes was eX1racted from the DESeq2 results. Principal Component Analysis (PCA) was conducted using the prcomp() function in RStudio. PC1 and PC2, which collectively accounted for the highest proportion of cumulative variance, were selected to generate a score plot for visualizing the distribution of sampies in the principal component space.

#### 2.10.4 Gene Ontology (GO) and Kyoto Encyclopedia of genes and genomes (KEGG) enrichment analysis

The target gene set obtained from the differential expression analysis was subjected to conversion from Gene Symbols to standardized Entrez IDs to facilitate subsequent GO and KEGG enrichment analyses. Significantly enriched terms were filtered using a threshold of adj.P < 0.05.

#### 2.10.5 Pseudotime analysis

The standardized expression matrix was imported into the R environment. Subsequently, trajectory inference was performed using the bulKPseudotime(). Specifically, highly variable genes were selected for trajectory construction. The program assigned a pseudotime value to each group along the trajectory, representing its progression state relative to the starting point To visualize the results, the expression dynamics of Key genes along the pseudotime axis were examined to validate the biological plausibility of the inferred trajectory.

### 2.11 Statistical analysis

Statistical analysis was performed using GraphPad Prism (version 9.0.0) or R (version 4.5.1). All results are presented as the mean ± standard deviation from at least three independent experiments. P values were calculated using Two-tailed student’s t-test or One-way analysis of variance (ANOVA), and P value < 0.05 was considered statistically significant.

## 3. Results

### 3.1 Establishment of Tet-On-PAX7 EPSCs

Leveraging the controllability of genetic modification in stem cells, we integrated a Tet-On system for inducible *PAX7* expression (Fig. 1A, S1A) into the pEPSC genome along with Id-Tomato gene using PiggyBac(Randolph et al., 2017) transposon technology. Successfully integrated positive clones were expected to express Id-Tomato. These positive clones were picked and expanded (Fig. 1B, top and botton). The expanded genetically modified stem cell colonies maintained typically dome-shape and stable red fluorescence expression (Fig. 1B, bottom). To assess whether genetic modification affected the pluripotency and Karyotype of pEPSCs, we performed AIKaline PhOsphatase (AP) staining (Fig. 1C), Karyotype analysis (Fig. 1D), RT.qPCR (Fig. 1F), and immunofluorescence staining for pluripotency marKers (SOX2, *OCT4,* and *NANOG)* (Fig. 1E) on Tet-On-PAX7 EPSCs. Results indicated that Tet-On-PAX7 EPSCs maintained a normal Karyotype, exhibited positive AP staining, and sustained stable expression of pluripotency marKers, comparable to WT EPSCs (Fig. S1B-O). Furthermore, we validated the inducible expression system of Tet-On-PAX7 EPSCs after adding 1 µg/ml DOX to the culture medium for 1 day. *PAX7* transcription levels were significantly upregulated, approximately 10,000-fold higher compared to the control group (Fig. 1G). These results demonstrate the successful establishment of a EPSC line capable of inducible *PAX7* expression while maintaining normal stem cell pluripotency and Karyotype.

**Figure 1.**
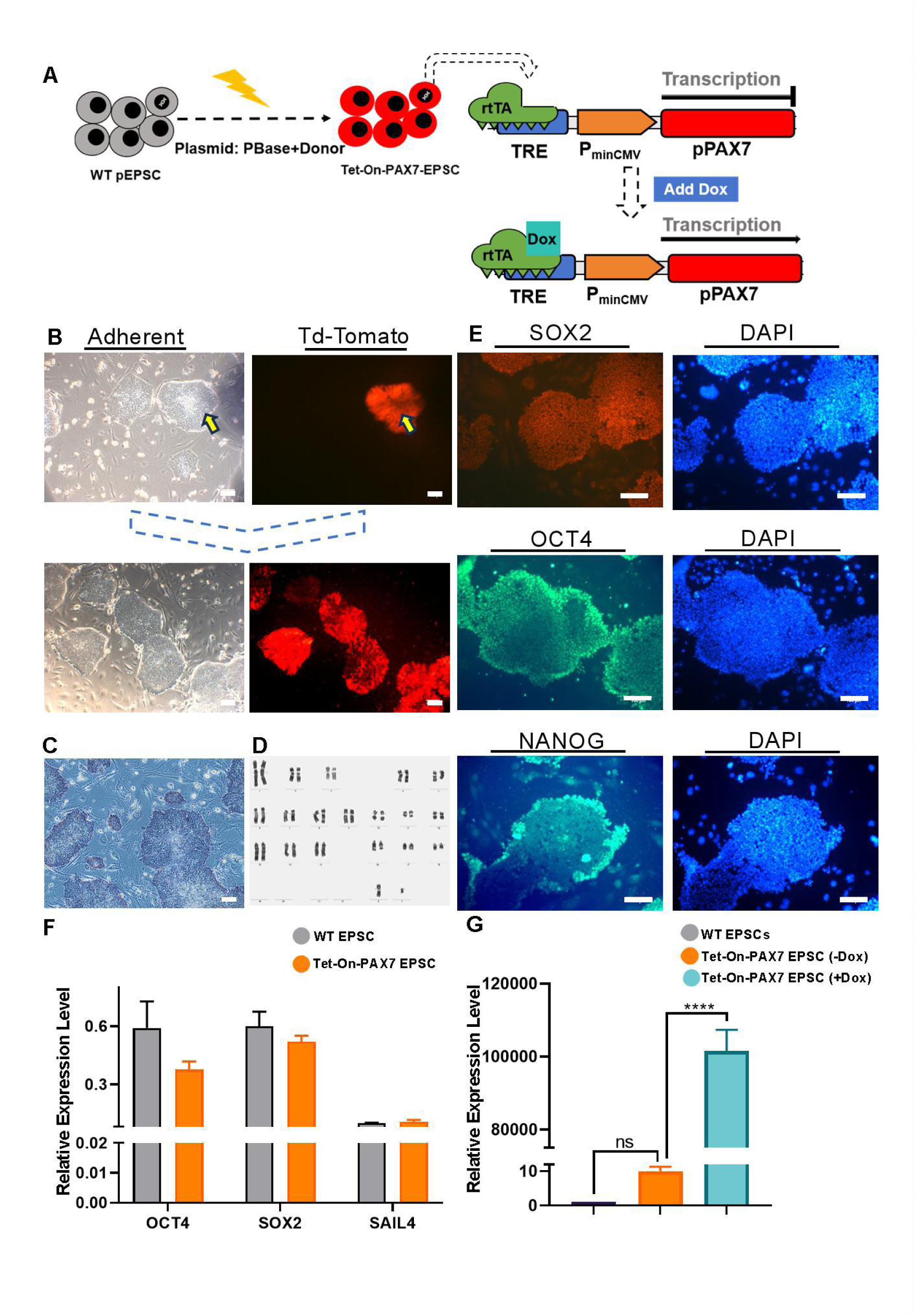
Genetically modified EPSCs maintain stable karyotypes and pluripotency with reliable Tet-On system functionality. **(A)** Schematic of pig Extended Pluripotent Stem Cells screening system and Tet-On inducible system: Successfully transfected cells express td-Tomato. In the absence of the inducer doxycycline (Dox), the expression of PAX7 is either silenced or maintained at a very low basal level. Upon Dox addition, Dox binds to the reverse tetracycline-controlled transactivator (rtTA), which subsequently activates the TRE promoter, specifically inducing high-level transcription of PAX7. **(B)** Td-Tomato-based selection and expansion of positive cell clones: Positive cell clones were selected based on red fluorescence signal, followed by passaging and expansion culture. **(C-F)** Pluripotency assessment and karyotyping analysis of Tet-On-PAX7 EPSCs: **(C)** Alkaline phosphatase staining, **(D)** karyotyping analysis, **(E)** Immunofluorescence staining, and **(F)** RT-qPCR analysis of pluripotency gene expression. **(G)** PAX7 expression level relative to the baseline after Dox induction. Note: n = 3 biological replicates. Statistical significance was calculated by Two-tailed student’s t-test. Data are presented as mean ± s.d. **** P < 0.0001. Scale bar, 100 μm.

### 3.2 The somite-like mesoderm cells established from Tet-On-PAX7 EPSCs exhibit high myogenic differentiation efficiency

We first differentiate Tet-On Pax7 EPSCs into somite-liKe mesoderm cells by treatment with ICS-ElnOe medium, which contains molecules promoting Wnt signaling activation (via CHIR99021, a GSK-3a/f3 inhibitor) and inhibiting TGF-13signaling (via SB-431542). After 12 days of culture in ICS-ElnOe, the resulted somite-liKe mesoderm cells lost the typical dome colony morphology of pEPSCs, exhibiting a relatively uniform monolayer appearance with predominantly spindle-shaped or irregularly polygonal cells, and distinct intercellular spaces (Fig. 2B). RT-qPCR confirmed that the expression levels of pluripotency factors such as *OCT4* and SOX2 were significantly downregulated, while somite-related marKers including *PAX3* and MEOX1(Conrad et al., 2023) were upregulated in somite-liKe mesoderm cells (Fig. 2C).

**Figure 2.**
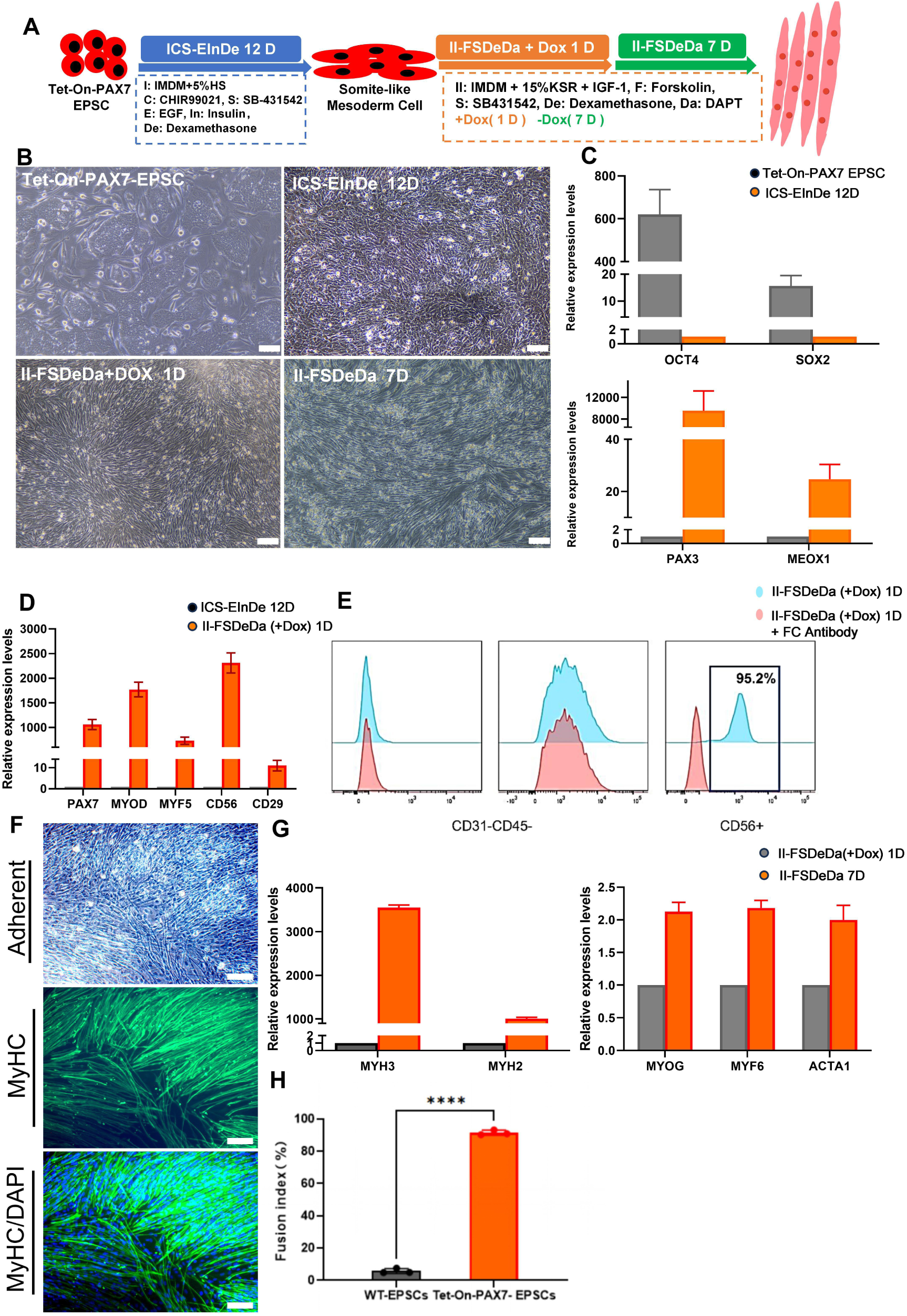
The somite-like mesoderm cells derived from Tet-On-PAX7 EPSCs exhibit high myogenic differentiation efficiency **(A)** The flowchart illustrates the process of differentiation from Tet-On-PAX7 EPSCs to myotube, along with the corresponding culture media and differentiation timelines. **(B)** Representative microscopic images showing the cellular morphology at different stages of differentiation. **(C)** Expression of pluripotency and somite-like mesoderm markers was detected by RT-qPCR: Following induction of differentiation in ICS-EInDe medium, the transcript levels of the pluripotency markers *OCT4* and *SOX2* decreased, while the expression of the somite-like mesoderm markers *PAX3* and *MEOX1* increased. **(D-E)** Muscle stem cell marker analysis: The II-FSDeDa medium containing Dox promoted the differentiation of somite-like mesoderm cells into muscle stem cells. Changes in the expression levels of muscle stem cell markers were detected by **(D)** RT-qPCR, and **(E)** flow cytometry analysis revealed a CD31-, CD45-, CD56+ phenotype. (F-G) Myogenic differentiation efficiency assessment: **(F)** MyHC immunofluorescence staining (green) was performed after 7 days of differentiation in II-FSDeDa medium, with DAPI staining the nuclei (blue). **(G)** The nuclear fusion efficiency was statistically analyzed. **(H)** RT-qPCR detection of myocyte and mature myotube markers: The expression levels of the myocyte markers *MYOG* and *MYF6*, and the myotube markers *MYH3*, *MYH2*, and *ACTA1*, were upregulated to varying degrees. Note: n = 3 biological replicates. Statistical significance was calculated by Two-tailed student’s t-test. Data are presented as mean ± s.d. **** P < 0.0001. Scale bar, 100 μm.

Subsequently, we designed a culture medium, 11-FSDeDa, based on the approach by Sridhar Selvaraj (Selvaraj et al., 2019), containing four small molecules that robustly promote myogenic differentiation and myotube fusion. The DOX was added to the medium to initiate the expression of *PAX7.* Through optimizing the induction concentration and duration of Oox treatment, we found that 1 µg/ml Oox applied for 1 day was sufficient to yield ideal myogenic differentiation outcomes (Fig. S2A-E). At this stage, the cells exhibited a relatively uniform elongated spindle morphology, along with an upregulation in the expression ofMuSC marKers *(PAX7, MYOD1, MYF5, CD56,* CO29) (Fig. 2D). Flow cytometry analysis showed that cells at this stage were *CD31^-^, CD45^-^* and *CD56^+^* (Fig. 2E), further confirming their Muse identity. After 7 days of culture in 11-FSDeDa without DOX, upregulated expression of myocyte and myotube marKers *(MYOG, MYF6, MYH3, MYH2, MYH4, ACTA1)* (Fig. 2G), and irregular contractions (Video S1) were observed, indicating differentiating progression toward mature myocyte and myotube differentiation. The immunofluorescence staining for myosin heavy chain (MyHC) on the differentiated cells (Fig. 2F) showed that a positive rate of approximately 92% was achieved (Fig. 2H). Serial time-point sampling analysis revealed that the temporal expression patterns of Key myogenic differentiation marKers recapitulated the *in-vivo* muscle development process (Fig. S2F). In summary, the somite-liKe mesoderm cells established from Tet-On-PAX7 EPSCs differentiated in ICS-ElnOe medium underwent highly efficient myogenic differentiation in 11-FSDeDa medium with 8 days.

### 3.3 Transcriptomic features across stages of myogenic differentiation

To elucidate the transcriptional heterogeneity of cells under different culture conditions, we performed RNA-seq analysis on four groups of cells: Tet-On-PAX7 EPSC, EPSC-Somite-liKe Mesoderm Cells (EPSC-SMC, ICS-ElnOe), EPSC-Muscle Stem Cell (EPSC-MuSC, 11-FSOeOa+Oox), and EPSC-Myocyte (EPSC-MC, 11-FSOeOa). Cross-stage differential expression analysis was conducted to construct a dynamic transcriptional landscape along the differentiation trajectory.

Overall, correlation analysis among samples showed that the intra-group correlation coefficients of all biological replicates were >-0.95 (Fig. 3A), confirming the robustness of technical reproducibility. AlthOugh the EPSC-MuSC group exhibited minor technical variation, all samples were strictly confined within the 95% confidence ellipse. Principal component analysis (PCA) further demonstrated that the cumulative contribution of PC1 and PC2 reached 92%, with distinct separation among groups in the 20 space (Fig. 3C), indicating statistically significant grouping. Each group contained a large number of uniquely differential expressed genes (unique OEGs), whereas the shared genes accounted for only 12.12% of the total, suggesting that cells at different differentiation stages possess clearly defined transcriptional features (Fig. 3B). Ternary plot analysis delineated a directed lineage trajectory from EPSC-SMC→EPSC-MuSC→EPSC-MC, with low transcriptional dispersion observed within populations (Fig. 3D). PC1 loading analysis further revealed that Tet-On-PAX7 EPSC highly expressed pluripotency-associated genes *(OCT4, NANOG),* while lineage-specific regulators *(PAX3,* PAX?) were progressively activated during differentiation, thereby driving myogenic commitment (Fig. 3E).

**Figure 3.**
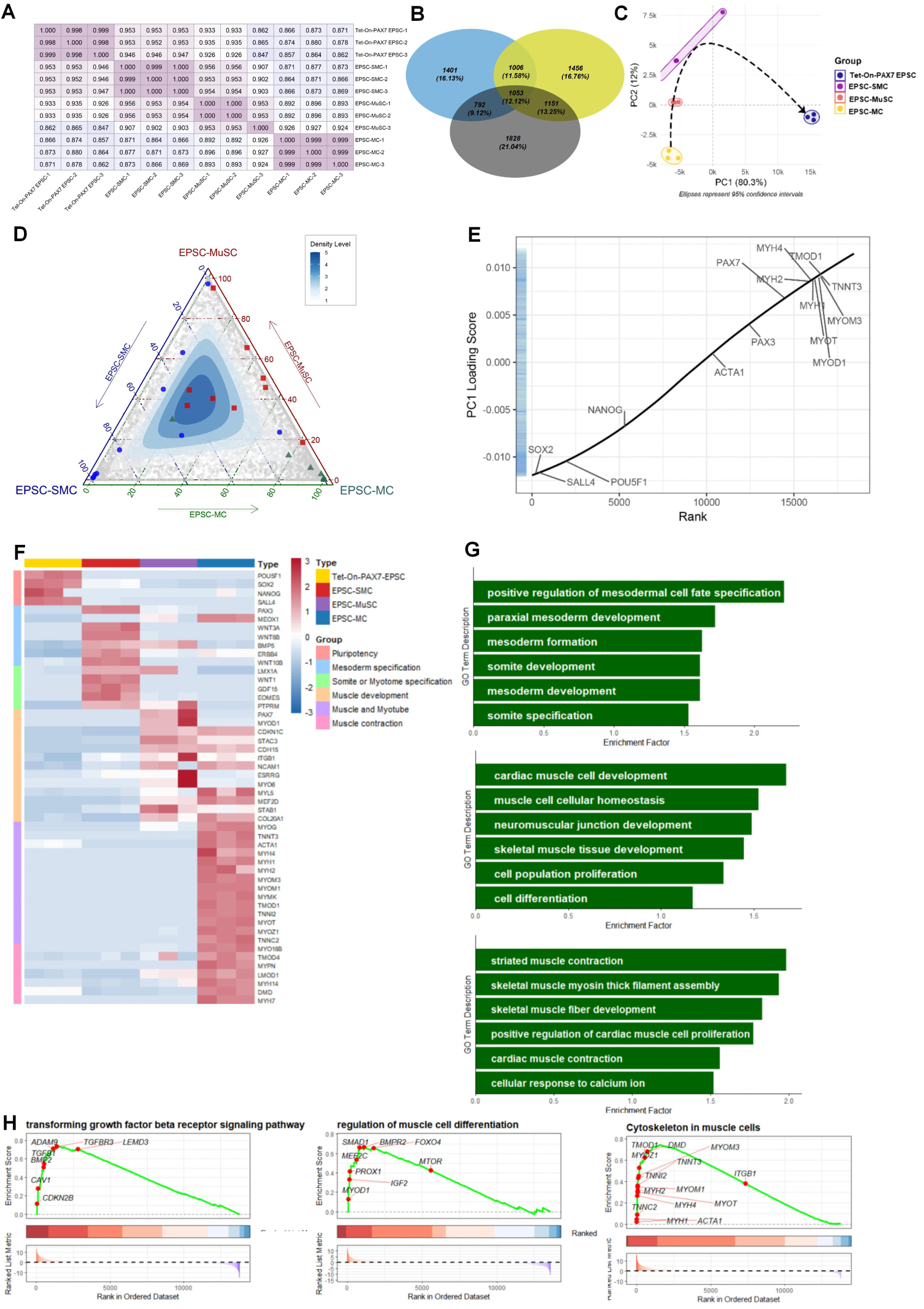
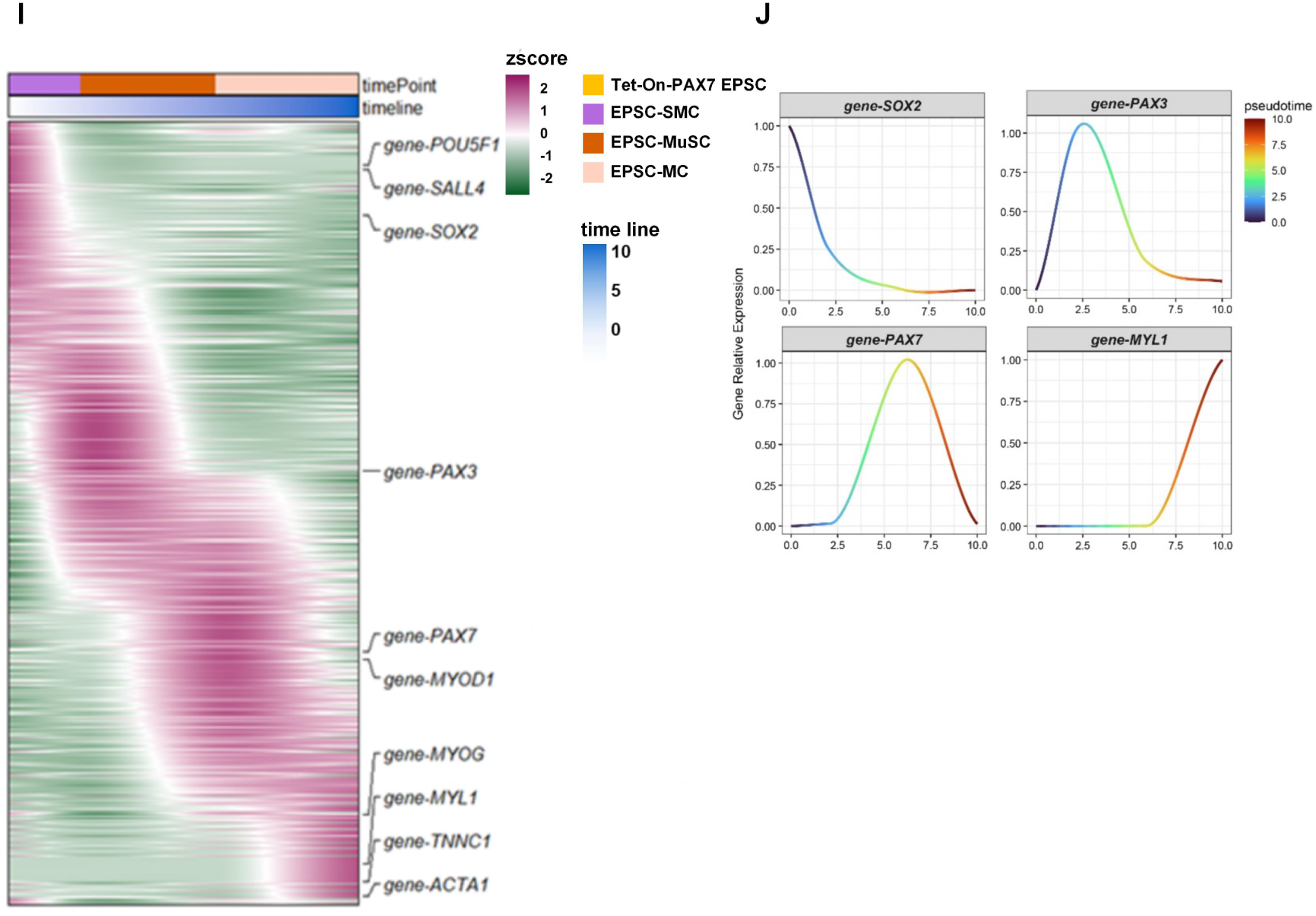
Transcriptomic features across stages of myogenic differentiation (A) Correlation heatmap among samples. (B) Venn diagram: Overlap of differentially expressed genes from pairwise comparisons between consecutive differentiation stages (Tet-On-PAX7-EPSC > EPSC-SMC >EPSC-MuSC >EPSC-MC). Tet-On-PAX7-EPSC vs EPSC-SMC (gray), EPSC-SMC vs EPSC-MuSC (blue), EPSC-MuSC vs EPSC-MC (yellow). (C) Principal component analysis (PCA): Transcriptomic differences among samples are displayed using the first two principal components (PC1 and PC2). Samples from the same group cluster within confidence ellipses, confirming the reliability of the grouping and suggesting that PC1 likely represents the primary developmental trajectory. (D) Ternary plot illustrating the transcriptomic features of three cell states (SMC, MuSC, MC). In the transcriptomic feature space defined by SMC, MuSC, MC key marker genes (blue circles: somite markers; red squares: muscle stem cell markers; green triangles: myocyte markers) are distributed along the anticipated path, validating the existence of the differentiation trajectory. (E) Distribution of gene expression along the PC1. (F) Heatmap of gene expression: The heatmap displays the expression patterns of genes from different functional modules across sample groups. Gene expression values are Z-score normalized (see the color bar on the right). Genes are clustered and grouped according to their functions. (G) GO pathway analysis and (H) GSEA results for relevant gene sets: Differentially expressed genes were derived from the comparisons Tet-On-PAX7-EPSC vs EPSC-SMC (top or left), EPSC-SMC vs EPSC-MuSC (middle), and EPSC-MuSC vs EPSC-MC (bottom or right). (I) Heatmap of the expression dynamics of key developmental genes along a pseudotemporal axis: The heatmap shows the dynamic changes in expression of key functional genes along the inferred pseudotemporal sequence (from left to right). Gene expression values are Z-score normalized. Each row represents a gene, and each column represents a cell’s position along the pseudotemporal axis. (J) Fitted curves of characteristic gene expression. Note: n = 3 biological replicates.

Analysis of OEGs between adjacent groups, followed by GO and KEGG enrichment, revealed stage-specific pathway activation. The OEGs between EPSC-SMC and Tet-On-PAX7 EPSC were enriched in the pathways associated with mesoderm specification and somitogenesis (Fig. S30, E). The differential expression genes between EPSC-MuSC and EPSC-SMC were concentrated in pathways related to cardiac and sKeletal muscle development (Fig. S4O, E). GSEA further demonstrated significant enrichment of “regulation of muscle differentiation” pathway in EPSC-MuSC, with leading-edge subsets dominated by canonical myogenic regulators *(MYOD1, MYF5).* Relative to EPSC-MuSC, EPSC-MC exhibited transcriptomic features more closely aligned with myocyte, characterized by elevated expression of genes involved in myofibril assembly and contractile function (Fig. 3F-H).

BulK RNA-seq pseudotime analysis captured the transcriptional dynamics underlying the transition from Tet-On-PAX7 EPSC to EPSC-MC (Fig. 3I, J). Heatmap visualization showed progressive silencing of pluripotency programs concurrent with activation of myogenic programs. Focusing on Key regulators (Fig. 3J), expression of the myosin gene *MYL1* and the myogenic factor *PAX?* increased continuously during differentiation, exhibiting lineage-specific activation patterns across different cell lines. These results confirm the orderly transcriptional activation of the myogenic differentiation program.

### 3.4 The somite-like mesoderm cells maintain high myogenic differentiation efficiency and genomic stability during long-term passaging

We next examined whether the somite-liKe mesoderm cells possess the abilities to maintain genomic stability and preserve differentiation efficiency even after long-term passaging. We performed long-term culture of the somite-liKe mesoderm cells (Fig. 4A) and plotted their proliferation curve. The cells at passage 40 had a more homogeneous morphology (Fig. 4B), a normal Karyotype (Fig. 4D), and no DNA damage confirmed by comet assays (Fig 4C). After 150 days of continuous culture, these cells underwent a 10^25^-fold expansion (Fig. 4E).

**Figure 4.**
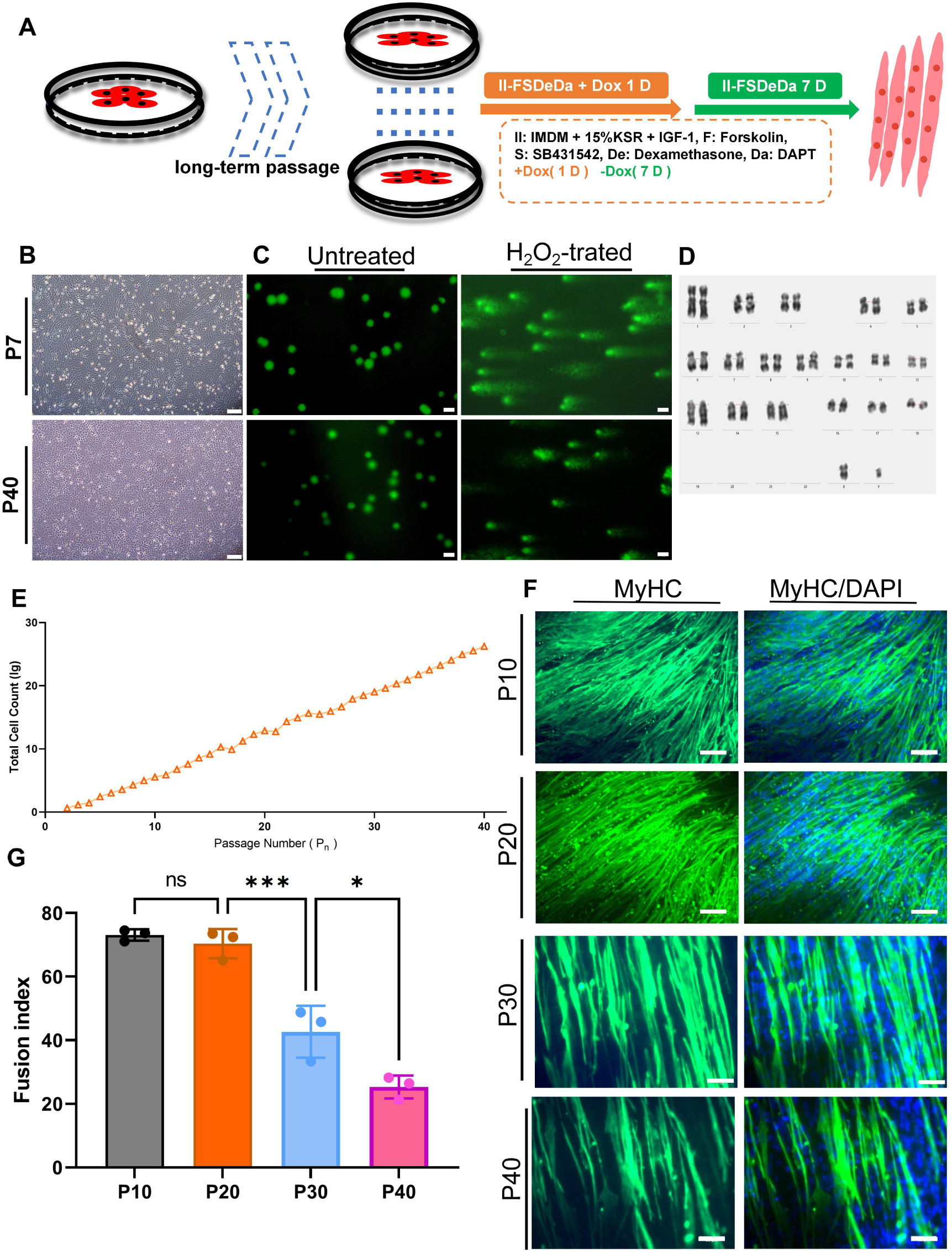
Genomic stability and myogenic differentiation efficiency of somite-like mesoderm cells during long-term passaging **(A)** Flowchart of long-term passage of somite-like mesoderm cells. **(B)** Morphology of somite-like mesoderm cells at passage 7 and passage 40 under microscopy. Scale bar, 40 μm. **(C)** Comet assay: Positive control treated with 3% H₂O₂. Compared to the untreated group (left), H₂O₂-treated cells (right) show significant comet tailing. Scale bar, 40 μm. **(D)** Karyotype analysis of passage 40 somite-like mesoderm cells. **(E)** Proliferation curve: Proliferation curve of somite-like mesoderm cells from passage 1 to passage 40. The Y-axis represents the total number of cells (lg), and the X-axis represents the number of cell passages. **(F)** MyHC immunofluorescence staining: MyHC staining (green) after myogenic differentiation of somite-like mesoderm cells at different passages (P10, P20, P30, P40). DAPI (blue) was used to stain cell nuclei, Scale bar, 100 μm or 50 μm. (G) Statistics of nuclear fusion efficiency at different passages. Note: n = 3 biological replicates. Statistical significance was calculated by One-Way ANOVA. Data are presented as mean ± s.d. ** P < 0.01.

To assess the impact of long-term passaging on myogenic capacity, we quantified the differentiation efficiency of somite-liKe mesoderm cells (at passages 10, 20, 30, and 40) by immunofluorescence staining for MyHC (Fig. 4F, G). The results demonstrated that somite-liKe mesoderm cells maintained a high differentiation efficiency of 70% up to passage 1Oto 20, and, cells at passage 30 and 40 still retained considerable myogenic potential, with efficiencies of 42.6% and 25%, respectively. These results indicate that somite-liKe mesoderm cells maintain genomic stability and robust myogenic differentiation capacity even after long-term passaging.

### 3.5 Somite-like mesoderm cells efficiently differentiated into myotubes on patterned and microcarrier scaffolds, as well as in anchorage-independent suspension culture

To validate the commercial potential of somite-liKe mesoderm cells, we cultivated them using patterned scaffolds, microcarrier scaffolds, and an anchor-free suspension system, which would confirm their suitability for large-scale production of porcine cultured meat.

The somite-liKe mesoderm cells were seeded in dishes and initially cultured in ordinary ICS-ElnOe medium. These cells then were cultured in 11-FSDeDa medium with 1 µg/ml Oox for a 1 day, and switched bacK to ordinary 11-FSDeDa medium for an additional day. The cells were collected and seeded onto patterned scaffolds (Fig. 5B). The cells effectively adhered to the patterned scaffolds. After cultured for 7 days, MyHC staining confirmed that they were able to differentiate into mature, textured myotube bundles (Fig. SC) in the patterned scaffolds. Spontaneous contraction of the myotube bundles on the scaffolds was observed (Video. S2).

**Figure 5.**
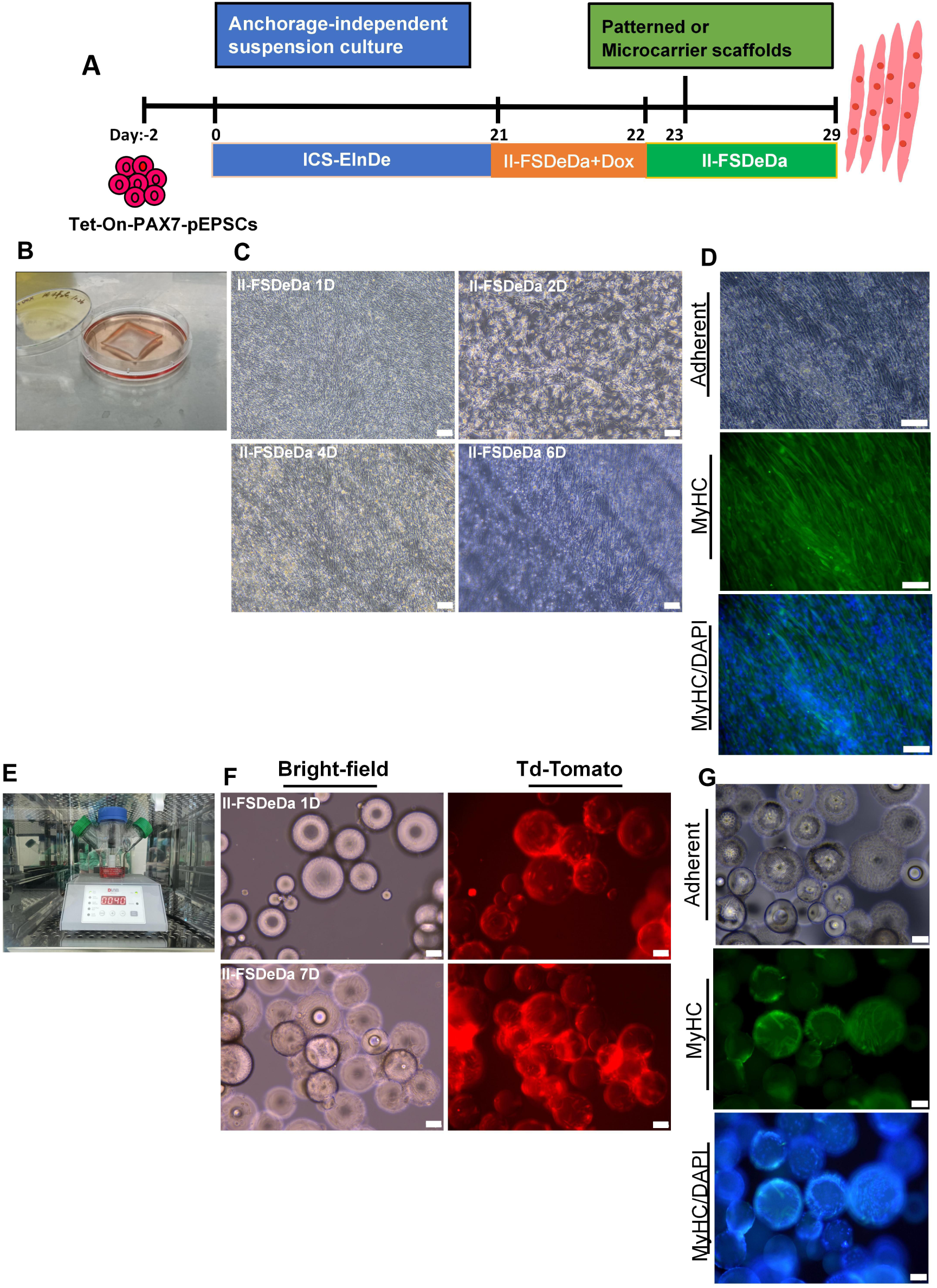
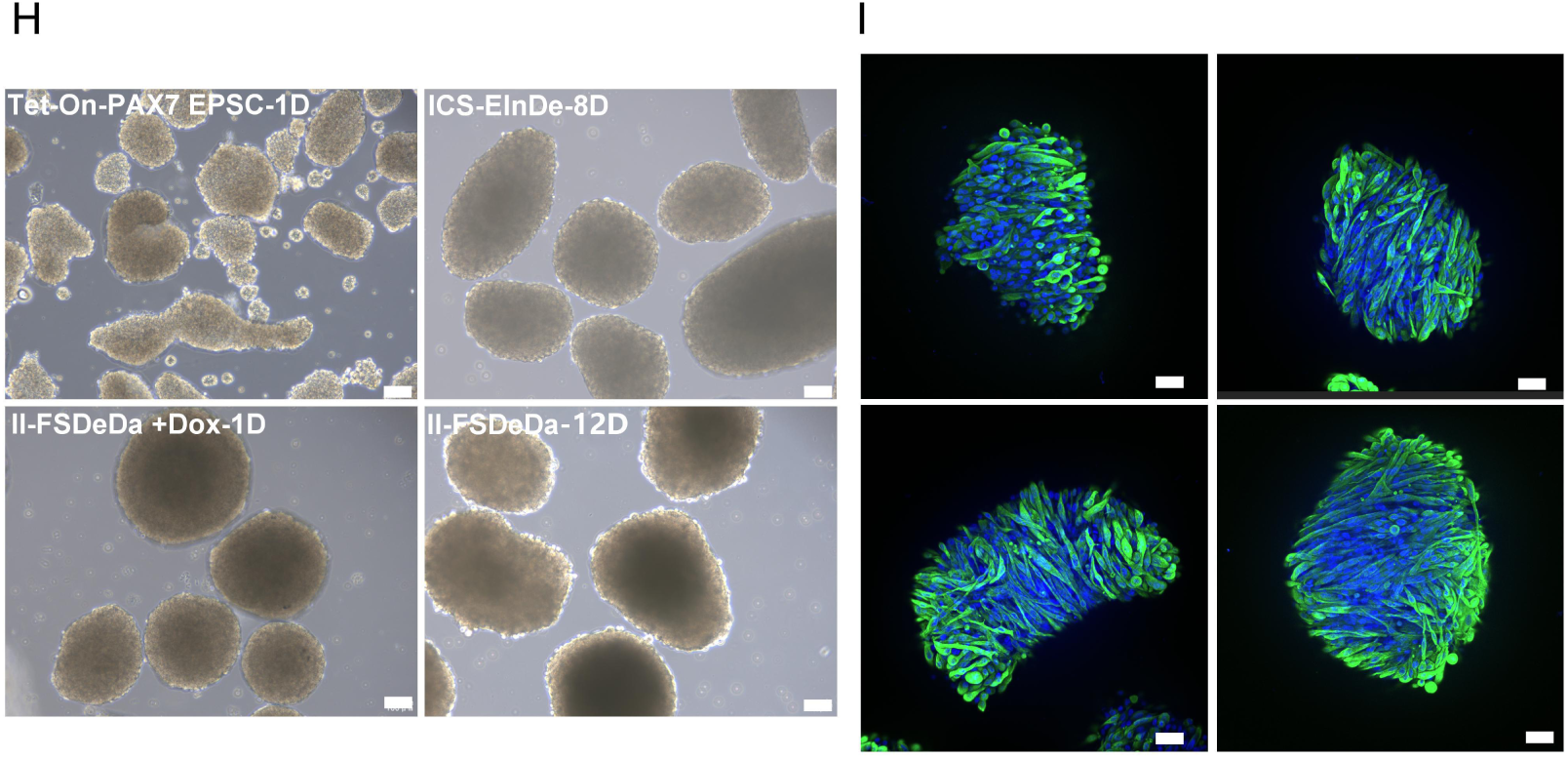
Somite-like mesoderm cells efficiently differentiated into myotubes in anchorage-independent suspension culture, as well as on patterned and microcarrier scaffolds **(A)** Flowchart illustrating myogenic differentiation of somite-like mesoderm cells in various culture system. Patterned scaffolds: **(B)** Myogenic differentiation of somite-like mesoderm cells in patterned scaffolds. **(C)** Temporal changes in cell morphology on patterned scaffolds from day 1 to day 6. **(D)** Efficiency of myogenic differentiation assessed by immunostaining for MyHC. Microsphere-based scaffolds: **(E)** Myogenic differentiation of somite-like mesoderm cells in microcarrier scaffolds under spinner flask system. **(F)** Cell adhesion on microcarriers in a spinner flask with II-FSDeDa medium. Bright-field and td-Tomato fluorescence images after 1 and 7 days of culture. **(G)** Efficiency of myogenic differentiation assessed by immunostaining for MyHC. Anchorage-independent suspension culture: **(H)** Temporal changes in cell morphology under various culture conditions. **(I)** Efficiency of myogenic differentiation assessed by immunostaining for MyHC. Note: n = 3 biological replicates. Scale bar, 100 μm.

Additionally, the cells were subjected to suspension culture within microcarrier scaffolds. In a spinner flask system, 2×10^6^ cells were seeded onto 20 mg of microcarriers (Fig. SE). Most cells with red fluorescence adhered to the microcarrier surfaces, and only very few suspension cells were observed in the medium (Fig. SF). After induction with Oox in 11-FSDeDa medium for one day, the medium was changed bacK to ordinary 11-FSDeDa medium and continuously cultured 7 days, the formation of myotubes was observed in this suspension system, confirmed MyHC staining (Fig. 5G).

Finally, we attempted to directly differentiate the cells into myotubes using an anchorage-independent suspension culture system. We observed that the somite-liKe mesoderm cells spontaneously aggregated into spheroids under spinner flasK culture conditions (Fig. SH). Then myogenic differentiation was induced using 11-FSOeOa medium with Oox for I day. After cultured for 7 days in 11-FSOeOa medium without Oox, the suspended spheroids exhibited spontaneous contraction (Video. S3), indicating the formation of myotubes. This was further confirmed by MyHC staining (Fig. 5I).

## 4. Discussion

Cultured meat technology represents an innovative approach to producing meat through *in-vitro* cell cultivation (Post et al., 2020). One of the major obstacles in cultured meat production is to obtain high-quality seed cells capable of both efficient differentiation and long-term passage. To overcome this challenge, we developed a novel seed cell: inducible PAX7-expressing somite-liKe mesodermal cells. This cell line has the high myogenic differentiation efficiency as Muse, as well as the sustained proliferation capacity as immortalized cells. Furthermore, these cells support robust myogenic differentiation across diverse culture environments, including 20 monolayers, 30 scaffold-based systems, and anchorage-independent suspension cultures, demonstrating their strong potential for scalable production and commercial application.

Previously, it was reported that treating human and mouse ESCs with the GSK3 inhibitor CHIR99021 followed by FGF2 and N2 supplements could result in approximately 90% of cells to achieve myogenic identity without any cell sorting. However, this small molecule-based approach tooK long time (more than 7 weeKs) to achieved myogenic identity from ESCs{Shelton et al., 2014). The recenuy proposed cultured meat production system{Zhu et al., 2023) utilizing pgEpiSCs as seed cells involves six different culture media, and requiring as long as 32-37 days plus N2 medium-mediated muscle cell maturation time (not displayed by the authors). Therefore, the differentiation process of pgEpiSCs based on small molecules into muscle represents a complex culture system with a lengthy timeline for production of cultured meat, which is an economical disadvantage.

Developmentally, PAX7, a crucial satellite cell marKer, is essential for driving somite differentiation toward the dermomyotome(Esteves de Lima and Relaix, 2021). In a previous study, DARABI et al(Oarabi and Perlingeiro, 2016) described a practical and efficient method for the derivation of sKeletal myogenic precursors from differentiating human pluripotent stem cells using controlled expression of PAX7. The myogenic precursors could be expanded exponentially and differentiated into myotubes *in-vitro* in a high efficiencies and shorter time, enabling researchers to use these cells for disease modeling as well as therapeutic purposes. Inspired by this finding, we used pEPSe as the initial stem cell source to produce cultured meat. The pEPSes is a type of formative-state pluripotent stem cells isolated from early blastocysts. They possess the ability to differentiate into both embryonic and extra-embryonic lineages(Gao et al., 2019; Ruan et al., 2024), being a more ideal stem cell source for their application in cultured meat production. We integrated the Tet-On inducible system for PAX7 expression into the pEPSes genome. The resulted Tet-On-PAX7 EPSes maintain normal pluripotency and can differentiate into somite-liKe mesoderm cells in IeS-ElnOe medium in 12 days. Most importantly, the somite-liKe mesoderm cells could differentiate into Muses in 11-FSDeDa medium merely in one day when treated with Oox to drive expression of PAX7 (Fig. 2D, E). The somite-liKe mesoderm cell-derived Muses could differentiate into muscle cells in 11-FSDeDa medium without Oox in 7 days, as evidenced by irregular contractions and the high expression of both fetal and adult muscle marKers, MYH3 and MYH2 (Fig. 2G)(Schiaffino et al., 2015). Integration of differential gene expression patterns along the Pe1 axis revealed the specific distribution of stage-specific marKer genes across distinct differentiation phases (Fig. 3E). This dynamic process was further corroborated by pseudotime analysis (Fig. 3I, J), collectively demonstrating the myogenic differentiation process of PAX7-expressing seed cells recapitulated *in-vivo* muscle development with a high efficiency.

In the process of production of cultured meat in our strategy, three types of stem cells are involved, pEPSes, somite-liKe mesoderm cells and Muses. Among them, pEPSCs have stronge proliferative and pluripotency, but they can be only used as initial seed cells due to their difficulty to maintain and lengthy process to differentiate into muscle as showed previously(Zhu et al., 2023). AlthOugh the differentiation of muscle stem cells into muscle fibers is relatively efficient(Post, 2014), duo to the difficulty in long-term passaging and the gradual loss of differentiation potential persistence typically reflected by impaired regenerative capacity and reduced myotube formation(BomKamp et al., 2022; Ding et al., 2017; Stout et al., 2023). Generally, the myogenic capacity of mammalian Muse can only be maintained for a limited number of passages *in-vitro,* around 1O passages(Xue et al., 2025). However, somite-liKe mesoderm cells could be passaged long-term (as long as 40 passages) cultured in ICS-ElnOe medium, with cell numbers expanding by 10^25^-fold (Fig. 3E), equivalent to approximately 83 population doublings, and still retaining a high myogenic differentiation capacity, 70%, 42.6% and 25% of resulted cells were myogenic cells for passages 20, 30 and 40, respectively (Fig.3F, G). The production of 1 Kg of cultured meat requires approximately 10^7^ cells(Zhu et al., 2022), and the proliferative capacity of a single somite-liKe mesoderm cell *in-vitro* far exceeds this demand. More notably, with only two induction media (ICS-ElnOe and 11-FSDeDa), our protocol generates myotubes from pEPSCs in approximately 20 days started form pEPSCs. This novel PAX7-based system thus offers a rapid, simplified, and efficient platform for advancing cultured meat research.

Biomaterial scaffolds are crucial for replicating the authentic sensory qualities of meat and facilitating the commercial production of cultured meat(Specht et al., 2018). Previously, three cultivation systems including 3O-printed patterned gelatin scaffolds, microcarrier scaffolds, and anchorage-independent suspension culture have been used for production of cultured meat. Among them, patterned scaffolds effectively replicate the structural and biochemical characteristics of natural muscle tissue(MacQueen et al., 2019), maKing them excellent materials for cultured meat production. For example, Li(Li et al., 2022) demonstrated that Chitosan-sodium alginate-collagen/gelatin 30 edible scaffolds supports cell growth and formation of appearance and texture similar to real meat. Microcarrier scaffolds (such as Seplife LX-MC-Dex1 microcarriers) offer a large surface area for cell adhesion and growth in stirred-tanK bioreactors(Acarregui et al., 2012; Ravi et al., 2015). The anchorage-independent suspension culture system, characterized by its operational simplicity, ease of scaling, and lower cost, represents a Key approach for commercial mass production by promoting efficient cell expansion and differentiation while overcoming the limitations of adherent culture. The U.S. FDA has accepted the conclusion that cultured meat products produced by GOOD Meat Company using suspension-adapted cell lines (FMT-SCF) in stirred-tanK bioreactors are safe for consumption(PasitKa et al., 2024). The somite-liKe mesoderm cells exhibited excellent adhesion capabilities on both patterned scaffolds and microcarrier scaffolds, also were able to spontaneously aggregate to form spheroids under anchorage-independent suspension culture conditions. And they successfully differentiated into myotubes with a high efficiency in all of the three scalable cultivation systems, validating their suitability for large-scale production of porcine meat.

In summary, a novel inducible PAX7-expressing somite-liKe mesoderm cell line was successfully established, demonstrating robust proliferative capacity alongside high myogenic differentiation potential to address a Key bottlenecK in cultured meat production. It exhibits excellent adaptability across various scalable cultivation systems, providing a versatile platform for high-density cell culture. However, future research should focus on refining culture medium components, such as eliminating animal-derived constituents from the ICS-ElnDe or optimizing the small molecule combinations in the 11-FSDeDa (Fig. S6), enhancing the myogenic differentiation capacity of high-passage cells, and investigating the nutritional profiles of cultured meat products, to facilitate the commercial application of this novel seed cell line.

## Compliance and ethics

The authors declare that they have no conflict of interest.

## Acknowledgements

This worKwas supported by National Key Research and Development Program of China (2022YFA1105400). Science and Technology Program of Guangzhou, China (2024803J1231). Science and Technology Planning Project of Guangdong Province, China (202381212060050, 202181212040016). We WOUid liKe to thanK the help provided by the Analysis and Testing Center of the Guangzhou Institutes of Biomedicine and Health, Chinese Academy of Sciences

## Author contributions

Cong Xia, Han Wu and Liangxue Lai conceived of the project. Cong Xia and Sheng Ye designed the experiments. Cong Xia and Sheng Ye completed the experimental design with help from Meichao Wang, Renquan Zhang and Xueliang Wang. Cong Xia analyzed the data with help from Hongliang Yu, Bingxiu Ma, Xiner Feng and Lijia Wen. Cong Xia drafted the manuscript. Cong Xia, Han Wu and Liangxue Lai reviewed and edited the manuscript. Yinghua Ye was responsible for the procurement of experimental supplies. All authors read and approved the final manuscript.

