## Supplementary Fig. S1-S6 for "Porcine extended pluripotent stem cell-derived somite-like mesoderm cells with Dox-driven PAX7 are robust seed cell resource for facilitating production of cultured meat"

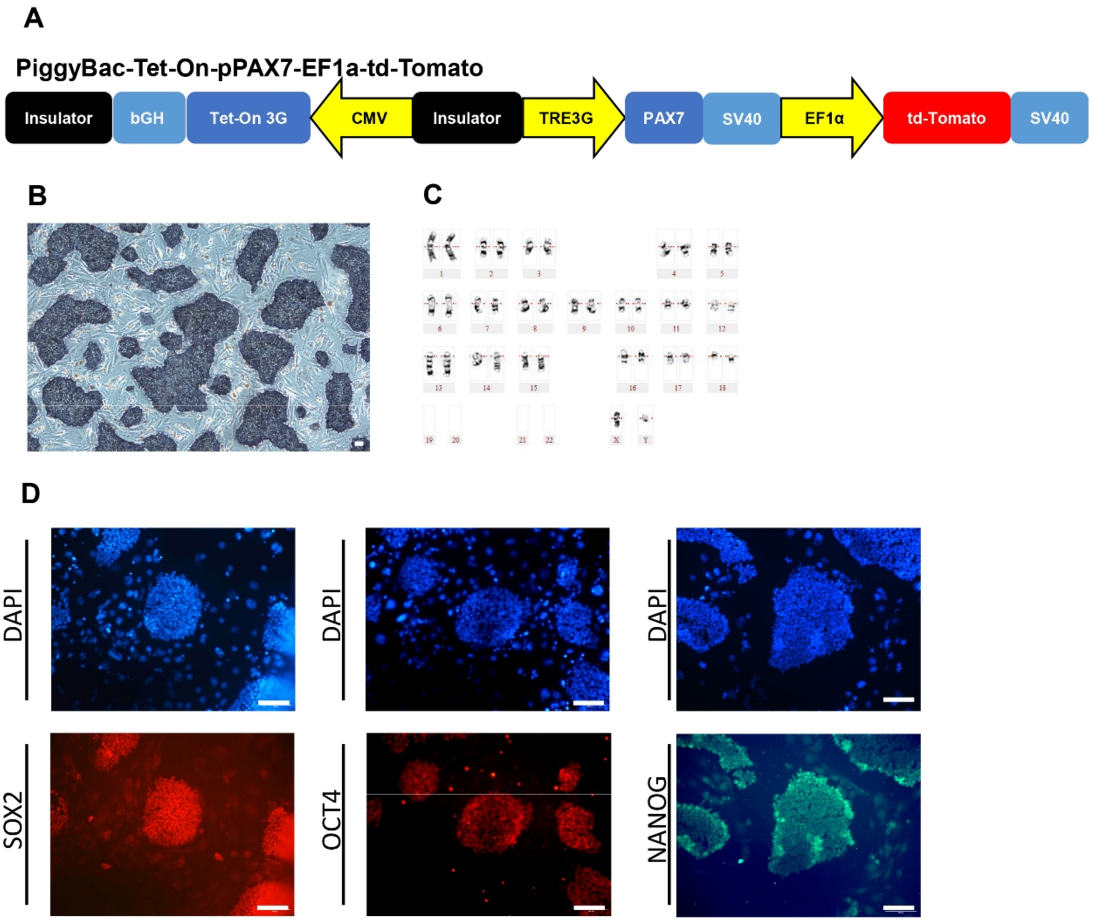


**Supplementary Figure.1 Vector Map and Wild-Type EPSC Pluripotency Identification**

(A) Schematic diagram of the Tet-On-PAX7-EF1a-td-Tomato plasmid; Pluripotency detection and karyotype analysis of WT-EPSCs: (B) AP staining, (C) karyotype analysis, and (D) pluripotency marker Immunofluorescence staining. Note: n = 3 biological replicates. Statistical significance was calculated by t-test. Data are presented as mean ± s.d. **** P < 0.0001.

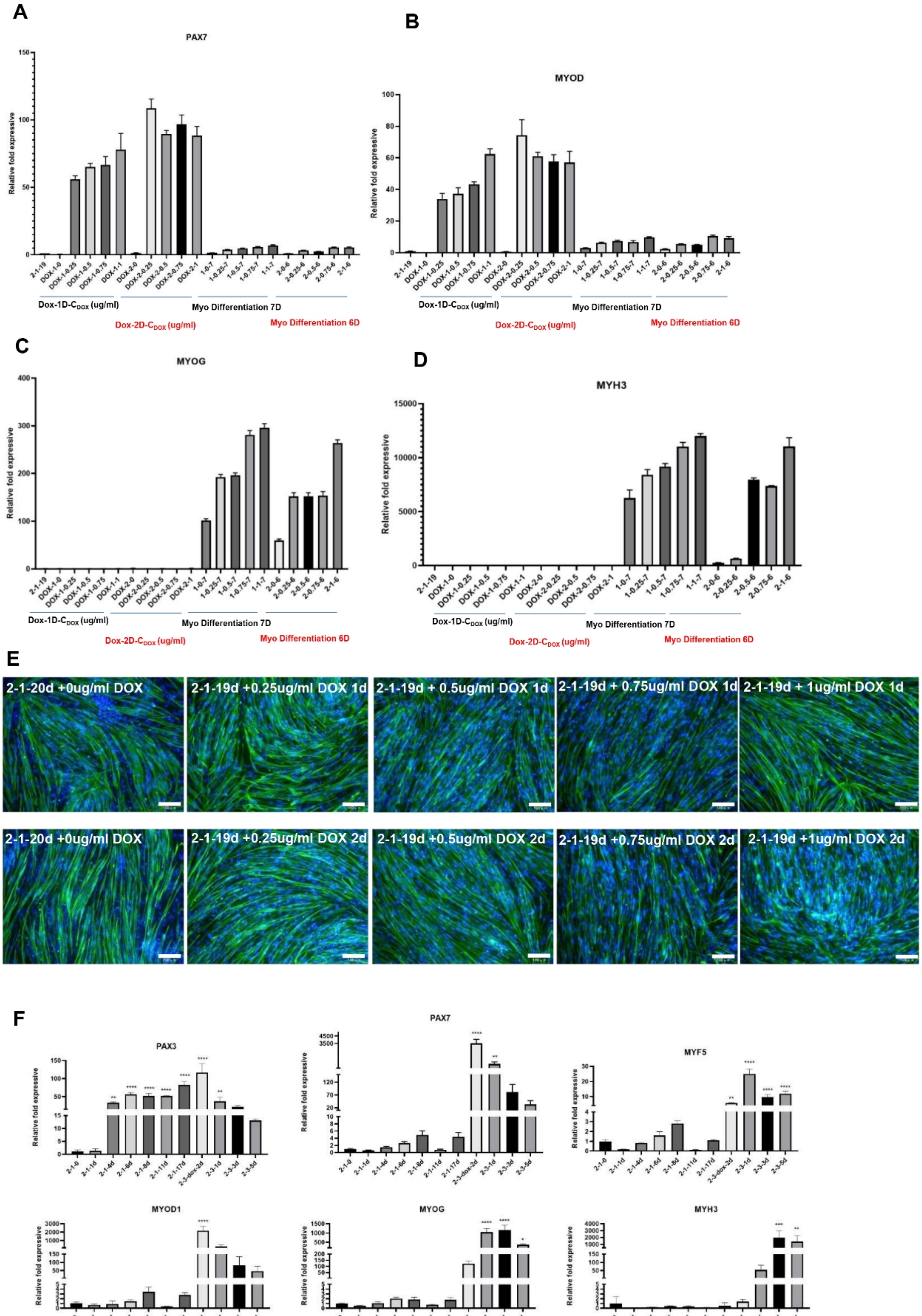
A
PAX7
Relative fold expressive
Dox-1D-C_DOX (ug/ml)
Myo Differentiation 7D
Dox-2D-C_DOX (ug/ml)
Myo Differentiation 6D
B
MYOD
C
MYOG
D
MYH3
E
2-1-20d +0ug/ml DOX
2-1-19d +0.25ug/ml DOX 1d
2-1-19d + 0.5ug/ml DOX 1d
2-1-19d + 0.75ug/ml DOX 1d
2-1-19d + 1ug/ml DOX 1d
2-1-20d +0ug/ml DOX
2-1-19d +0.25ug/ml DOX 2d
2-1-19d +0.5ug/ml DOX 2d
2-1-19d +0.75ug/ml DOX 2d
2-1-19d +1ug/ml DOX 2d
F
PAX3
PAX7
MYF5
MYOD1
MYOG
MYH3

**Supplementary Figure.2 Optimization of Dox Concentration and Treatment Duration for Myogenic Differentiation**

(A-D) Transcriptional levels of key myogenic genes assessed by RT-qPCR under varying Dox concentrations (0, 0.25, 0.5, 0.75, 1.0 μg/mL) and treatment durations (1 or 2 days). Samples were collected at two time points: after 1 or 2 days of Dox induction and at the end of the 7 or 6-day differentiation protocol (total differentiation time: 8 days). Genes analyzed: (A) *PAX7*, (B) *MYOD*, (C) *MYOG*, (D) *MYH3*. (E) Myogenic differentiation efficiency evaluated by MyHC immunostaining across treatment groups. Control: Precursor seed cells maintained in ICS-ElnDe (abbreviated as 2-1) medium for 19 days without Dox induction (denoted as 2-1-19). (F) Expression profiling of stage-specific markers via RT-qPCR: somite-like mesoderm (*PAX3*), muscle stem cell (*PAX7*, *MYOD*), and myocyte/myotube (*MYF5*, *MYOG*, *MYH3*).

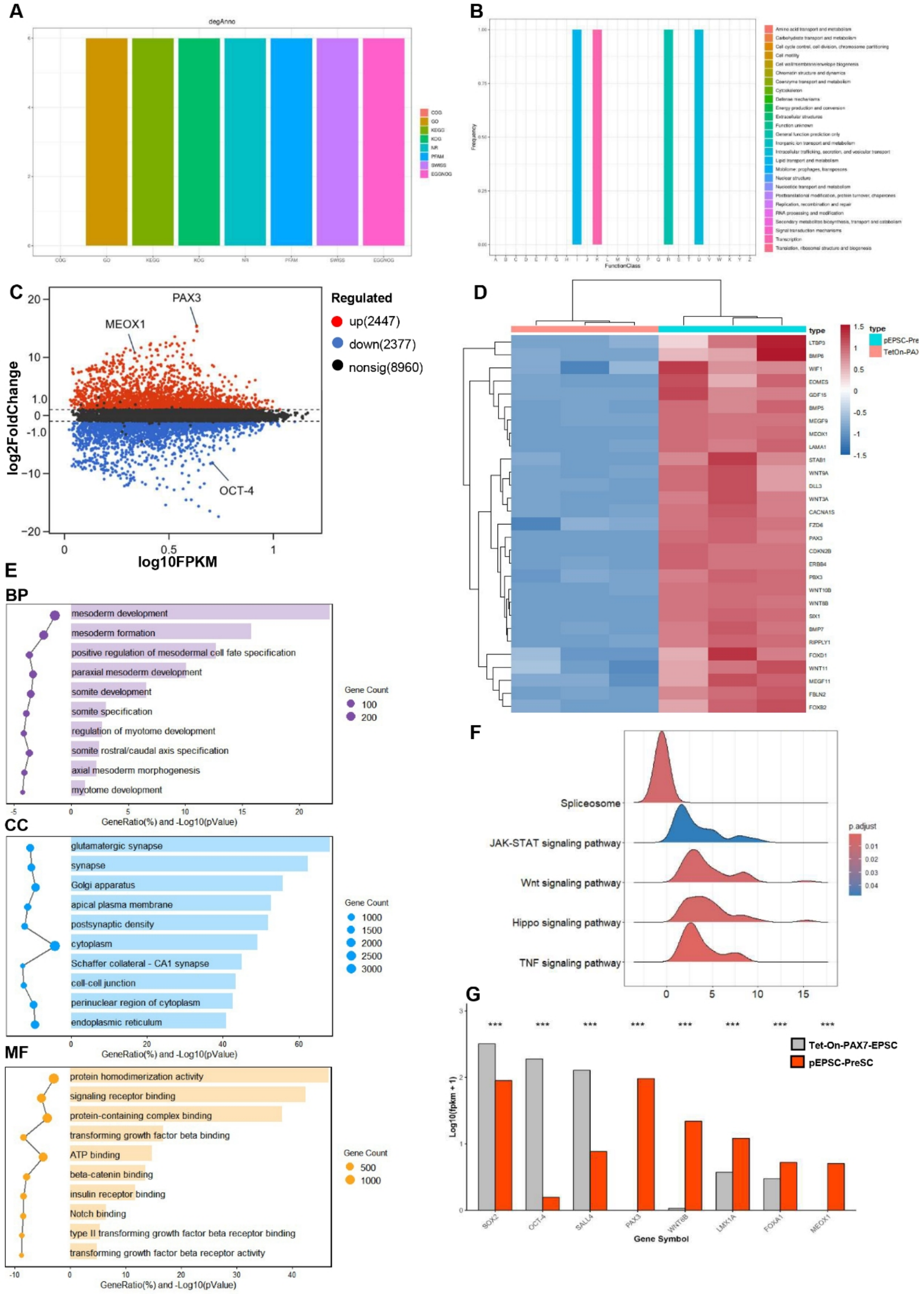
A
degAnno
COG
GO
KEGG
KOG
NR
PFAM
SWISS
EGGNOG
B
Frequency
FunctionClass
C
PAX3
MEOX1
OCT-4
log2FoldChange
log10FPKM
Regulated
up(2447)
down(2377)
nonsig(8960)
D
type
pEPSC-Pre
TetOn-PAX
LTBP3
BMP6
WIF1
EOMES
GDF15
BMP5
MEGF9
MEOX1
LAMA1
STAB1
WNT9A
DLL3
WNT3A
CACNA1S
FZD6
PAX3
CDKN2B
ERBB4
PBX3
WNT10B
WNT8B
SIX1
BMP7
RIPPLY1
FOXD1
WNT11
MEGF11
FBLN2
FOXB2
E
BP
mesoderm development
mesoderm formation
positive regulation of mesodermal cell fate specification
paraxial mesoderm development
somite development
somite specification
regulation of myotome development
somite rostral/caudal axis specification
axial mesoderm morphogenesis
myotome development
GeneRatio(%) and -Log10(pValue)
Gene Count
CC
glutamatergic synapse
synapse
Golgi apparatus
apical plasma membrane
postsynaptic density
cytoplasm
Schaffer collateral - CA1 synapse
cell-cell junction
perinuclear region of cytoplasm
endoplasmic reticulum
MF
protein homodimerization activity
signaling receptor binding
protein-containing complex binding
transforming growth factor beta binding
ATP binding
beta-catenin binding
insulin receptor binding
Notch binding
type II transforming growth factor beta receptor binding
transforming growth factor beta receptor activity
F
Spliceosome
JAK-STAT signaling pathway
Wnt signaling pathway
Hippo signaling pathway
TNF signaling pathway
p.adjust
G
Tet-On-PAX7-EPSC
pEPSC-PreSC
Log10(fpkm + 1)
SOX2
OCT-4
SALL4
PAX3
WNT8B
LMX1A
FOXA1
MEOX1
Gene Symbol

**Supplementary Figure.3 Clustering Analysis of Differentially Expressed Genes during Differentiation from Tet-On-PAX7-EPSC to pEPSC-PreSC**

(A) Differential Gene Annotation (degAnno) : The number of differentially expressed genes annotated in the database displayed. (B) Differential Gene Annotation (KOG) : The number of differentially expressed genes annotated in the KOG (Eukaryotic Orthologous Groups) database. (C) MA-Plot of Differential Gene Expression Analysis : The x-axis represents the average expression level of genes across all samples, while the y-axis represents the log2 fold change in gene expression. Points in the plot represent individual genes: red points indicate significantly upregulated differentially expressed genes (|log2FoldChange| > 1 and p.adjust < 0.05), blue points indicate significantly downregulated differentially expressed genes, and gray points indicate genes with no significant difference. (D) Heatmap of Differential Gene Clustering . (E) GO Enrichment Analysis of Differentially Expressed Genes : Displaying significantly enriched Biological Processes (BP), Cellular Components (CC), and Molecular Functions (MF). The bubble plot on the left has the x-axis representing the scaled gene ratio (GeneRatio), and the bubble size represents the number of genes enriched in the corresponding pathway. (F) Ridgeline Plot of Specific KEGG Pathway Enrichment Based on GSEA : The plot shows the gene distribution of six specific significantly enriched pathways. The x-axis represents the Normalized Enrichment Score (NES), and the y-axis lists the target pathway names. The ridge shape distribution for each pathway represents the NES density of the core enriched genes within that pathway. The ridge height reflects the concentration of gene distribution in the ranked list, and the fill color depth represents p.adjust. (G) Expression Levels of Selected Candidate Genes.

Note: n = 3. Statistical significance was calculated by t-test. Data are presented as mean ± s.d. * $p < 0.05$, ** $p < 0.01$, *** $p < 0.001$.

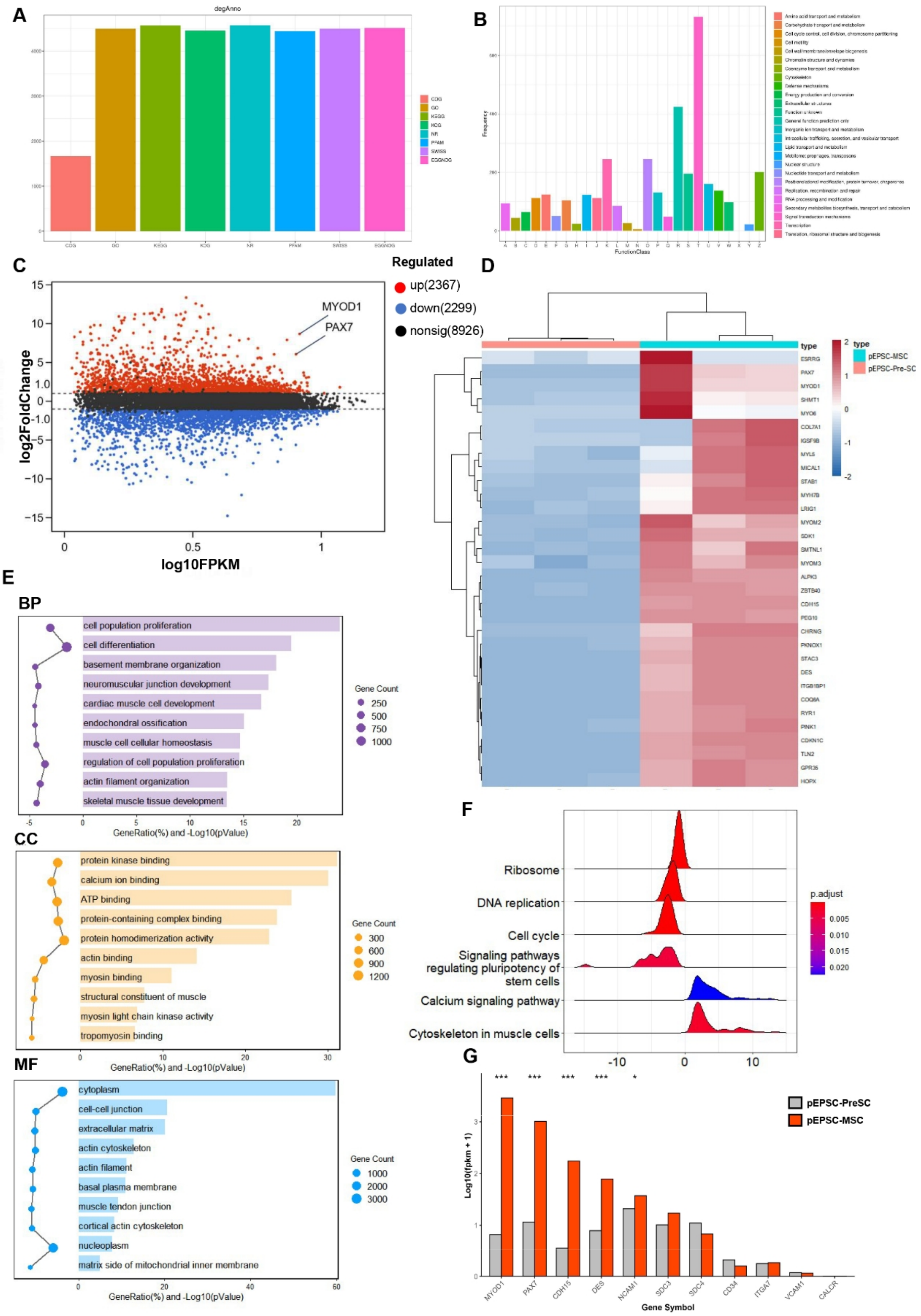
A
degAnno
COG
GO
KEGG
KOG
NR
PFAM
SWISS
EGGNOG
B
Frequency
FunctionClass
C
Regulated
up(2367)
down(2299)
nonsig(8926)
MYOD1
PAX7
log2FoldChange
log10FPKM
D
type
pEPSC-MSC
pEPSC-Pre-SC
ESRRG
PAX7
MYOD1
SHMT1
MYO6
COL7A1
IGSF9B
MYL5
MICAL1
STAB1
MYH7B
LRIG1
MYOM2
SDK1
SMTNL1
MYOM3
ALPK3
ZBTB40
CDH15
PEG10
CHRNG
PKNOX1
STAC3
DES
ITGB1BP1
COQ8A
RYR1
PINK1
CDKN1C
TLN2
GPR35
HOPX
E
BP
cell population proliferation
cell differentiation
basement membrane organization
neuromuscular junction development
cardiac muscle cell development
endochondral ossification
muscle cell cellular homeostasis
regulation of cell population proliferation
actin filament organization
skeletal muscle tissue development
Gene Count
250
500
750
1000
GeneRatio(%) and -Log10(pValue)
CC
protein kinase binding
calcium ion binding
ATP binding
protein-containing complex binding
protein homodimerization activity
actin binding
myosin binding
structural constituent of muscle
myosin light chain kinase activity
tropomyosin binding
Gene Count
300
600
900
1200
GeneRatio(%) and -Log10(pValue)
MF
cytoplasm
cell-cell junction
extracellular matrix
actin cytoskeleton
actin filament
basal plasma membrane
muscle tendon junction
cortical actin cytoskeleton
nucleoplasm
matrix side of mitochondrial inner membrane
Gene Count
1000
2000
3000
GeneRatio(%) and -Log10(pValue)
F
Ribosome
DNA replication
Cell cycle
Signaling pathways regulating pluripotency of stem cells
Calcium signaling pathway
Cytoskeleton in muscle cells
p.adjust
0.005
0.010
0.015
0.020
G
pEPSC-PreSC
pEPSC-MSC
Log10(fpkm + 1)
MYOD1
PAX7
CDH15
DES
NCAM1
SDC3
SDC4
CD34
ITGA7
VCAM1
CALCR
Gene Symbol

**Supplementary Figure.4 Clustering Analysis of Differentially Expressed Genes during Differentiation from pEPSC-PreSC to pEPSC-MSC**

(A) DegAnno : The number of differentially expressed genes annotated in the database displayed. (B) Differential Gene Annotation (KOG). (C) MA-Plot of Differential Gene Expression Analysis. (D) Heatmap of Differential Gene Clustering . (E) GO Enrichment Analysis of Differentially Expressed Genes. (F) Ridgeline Plot of Specific KEGG Pathway Enrichment Based on GSEA . (G) Expression Levels of Selected Candidate Genes.

Note: n = 3. Statistical significance was calculated by t-test. Data are presented as mean ± s.d. * $p < 0.05$, ** $p < 0.01$, *** $p < 0.001$.

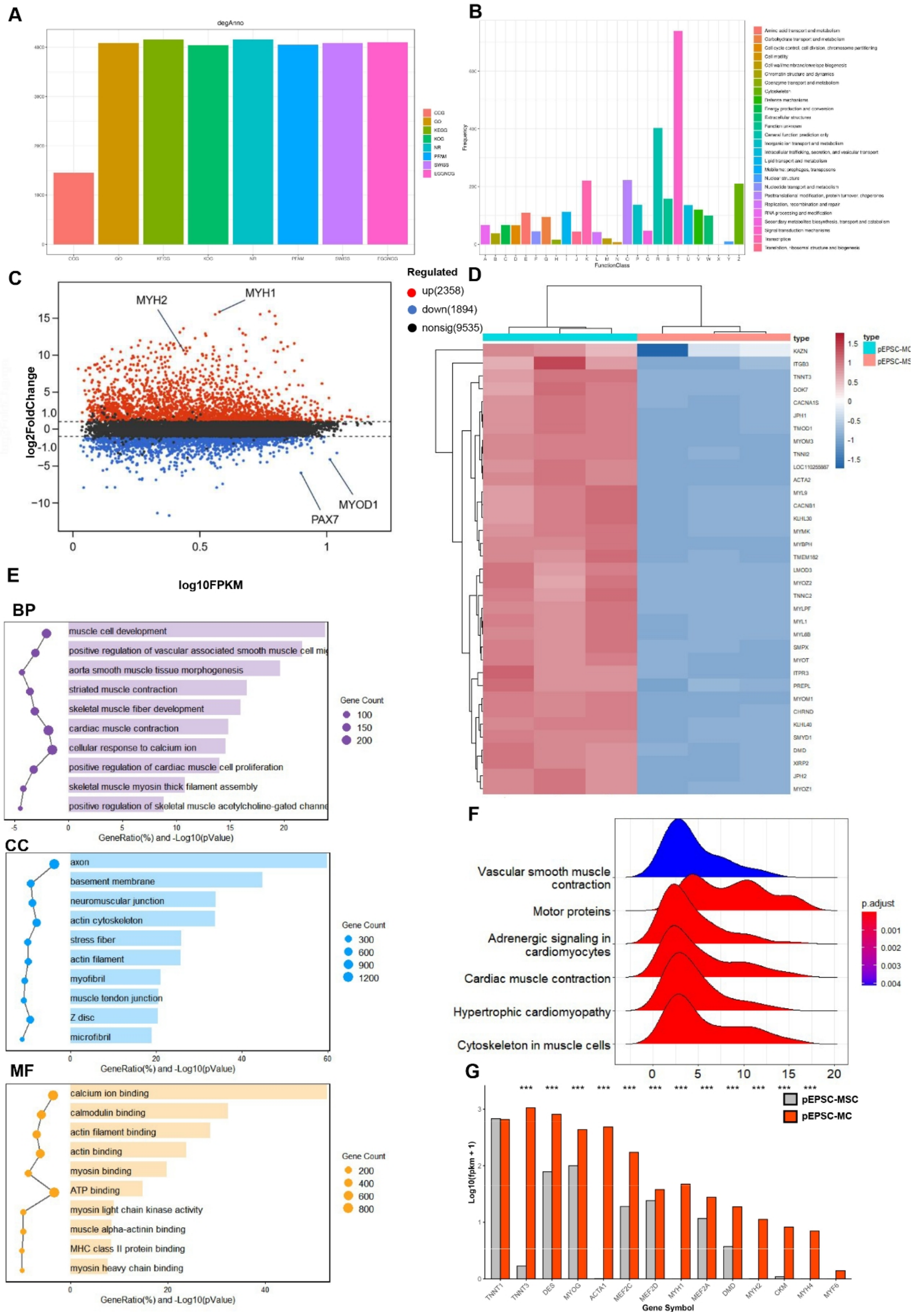

A
degAnno
COG
GO
KEGG
KOG
NR
PFAM
SWISS
EGGNOG
B
Frequency
FunctionClass
C
MYH2
MYH1
MYOD1
PAX7
log2FoldChange
log10FPKM
Regulated
up(2358)
down(1894)
nonsig(9535)
D
type
pEPSC-MC
pEPSC-MSC
KAZN
ITGB3
TNNT3
DOK7
CACNA1S
JPH1
TMOD1
MYOM3
TNNI2
LOC110255887
ACTA2
MYL9
CACNB1
KLHL30
MYMK
MYBPH
TMEM182
LMOD3
MYOZ2
TNNC2
MYLPF
MYL1
MYL6B
SMPX
MYOT
ITPR3
PREPL
MYOM1
CHRND
KLHL40
SMYD1
DMD
XIRP2
JPH2
MYOZ1
E
BP
muscle cell development
positive regulation of vascular associated smooth muscle cell mi
aorta smooth muscle tissue morphogenesis
striated muscle contraction
skeletal muscle fiber development
cardiac muscle contraction
cellular response to calcium ion
positive regulation of cardiac muscle cell proliferation
skeletal muscle myosin thick filament assembly
positive regulation of skeletal muscle acetylcholine-gated channe
GeneRatio(%) and -Log10(pValue)
Gene Count
100
150
200
CC
axon
basement membrane
neuromuscular junction
actin cytoskeleton
stress fiber
actin filament
myofibril
muscle tendon junction
Z disc
microfibril
GeneRatio(%) and -Log10(pValue)
Gene Count
300
600
900
1200
MF
calcium ion binding
calmodulin binding
actin filament binding
actin binding
myosin binding
ATP binding
myosin light chain kinase activity
muscle alpha-actinin binding
MHC class II protein binding
myosin heavy chain binding
GeneRatio(%) and -Log10(pValue)
Gene Count
200
400
600
800
F
Vascular smooth muscle contraction
Motor proteins
Adrenergic signaling in cardiomyocytes
Cardiac muscle contraction
Hypertrophic cardiomyopathy
Cytoskeleton in muscle cells
p.adjust
0.001
0.002
0.003
0.004
G
pEPSC-MSC
pEPSC-MC
Log10(fpkm + 1)
Gene Symbol
TNNT1
TNNT3
DES
MYOG
ACTA1
MEF2C
MEF2D
MYH1
MEF2A
DMD
MYH2
CKM
MYH4
MYF6

**Supplementary Figure.5 Clustering Analysis of Differentially Expressed Genes during Differentiation from pEPSC-MSC to pEPSC-MSC**

(A) DegAnno : The number of differentially expressed genes annotated in the database displayed. (B) Differential Gene Annotation (KOG). (C) MA-Plot of Differential Gene Expression Analysis. (D) Heatmap of Differential Gene Clustering . (E) GO Enrichment Analysis of Differentially Expressed Genes. (F) Ridgeline Plot of Specific KEGG Pathway Enrichment Based on GSEA . (G) Expression Levels of Selected Candidate Genes.

Note: n = 3. Statistical significance was calculated by t-test. Data are presented as mean ± s.d. * $p < 0.05$, ** $p < 0.01$, *** $p < 0.001$.

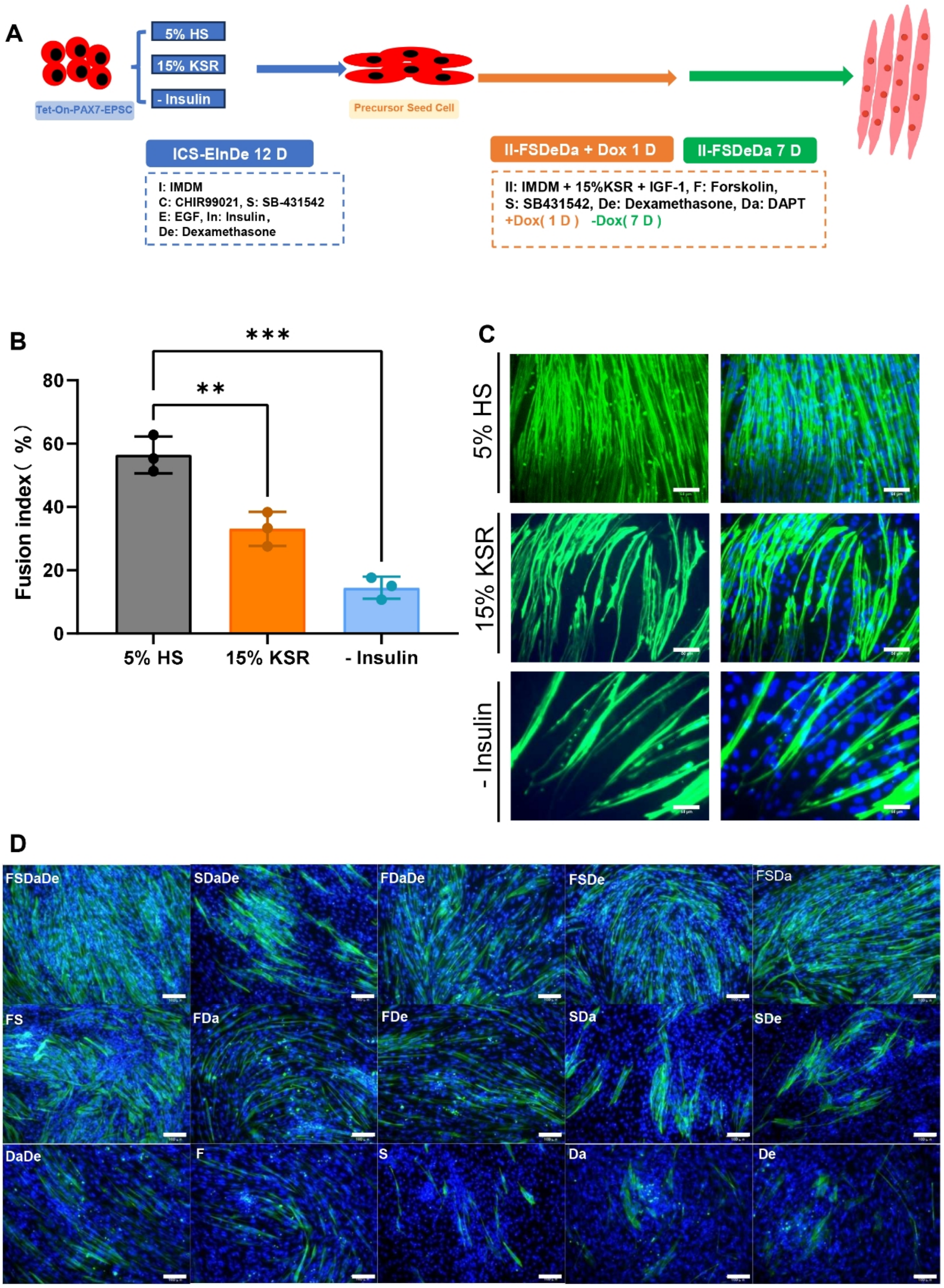
A
5% HS
15% KSR
- Insulin
Tet-On-PAX7-EPSC
Precursor Seed Cell
ICS-EInDe 12 D
I: IMDM
C: CHIR99021, S: SB-431542
E: EGF, In: Insulin,
De: Dexamethasone
II-FSDeDa + Dox 1 D
II-FSDeDa 7 D
II: IMDM + 15%KSR + IGF-1, F: Forskolin,
S: SB431542, De: Dexamethasone, Da: DAPT
+Dox( 1 D ) -Dox( 7 D )
B
***
**
Fusion index ( % )
80
60
40
20
0
5% HS
15% KSR
- Insulin
C
5% HS
15% KSR
- Insulin
D
FSDaDe
SDaDe
FDaDe
FSDe
FSDa
FS
FDa
FDe
SDa
SDe
DaDe
F
S
Da
De

**Supplementary Figure.6 Optimization of culture medium formulations, involving the elimination of animal-derived components from the ICS-EInD medium and the optimization of small molecule combinations in II-FSDeDa**

(A) Schematic of the ICS-EInDe optimization process: 5% HS was replaced with 15% KSR, or insulin was removed, followed by subsequent differentiation procedures. The ICS-EInD medium containing 5% HS served as the reference. (B) Comparison of differentiation outcomes among the three medium groups via MyHC immunofluorescence staining. (C) Statistics of nuclear fusion efficiency. (D) Assessment of myogenic differentiation efficiency in II-FSDeDa under different small molecule combinations by MyHC immunofluorescence staining.

**Supplementary Table.1 Primers used in RT-qPCR**

| Name | Sequence |
|---|---|
| pig-GAPDH-qPCR-F* | GGGCATGAACCATGAGAAGT |
| pig-GAPDH-qPCR-R* | GTCTTCTGGGTGGCAGTGAT |
| pig-SOX2-qPCR-F* | AAGAGAACCCCAAGATGCACAACT |
| pig-SOX2-qPCR-R* | GCTTGGCCTCGTCGATGAAC |
| pig-OCT4-qPCR-F* | GCAAACGATCAAGCAGTGA |
| pig-OCT4-qPCR-R* | GGTGACAGACACCGAGGGA |
| pig-NANOG-qPCR-F* | TTCCTTCCTCCATGGATCTG |
| pig-NANOG-qPCR-R* | ATCTGCTGGAGGCTGAGGTA |
| pig-pPAX7-qPCR-F | TCCTCTGGAAGTGTCCACCC |
| pig-pPAX7-qPCR-R | GCAGGGGTCGCCCATTGATG |
| pig-PAX3-qPCR-F | GCAGTATGGACAAAGTGCCT |
| pig-PAX3-qPCR-R | CAGGGCCAGTTTAAGCTCCA |
| pig-MEOX1-qPCR-F | AAAGGAGAGTTCAGGTCAAAGT |
| pig-MEOX1-qPCR-R | GGAGATGGGCTGACCGC |
| pig-MYOD-qPCR-F | TTCCGACGGCATGATGGATT |
| pig-MYOD-qPCR-R | TCGCTGTAATAGGTGCCGTC |
| pig-MYF5-qPCR-F | CCGACGGCATGCCTGAAT |
| pig-MYF5-qPCR-R | TTGGTACATCCGGACAGTAGA |
| pig-CD31-qPCR-F1 | CGAGGTCTGGGAACAAAGGG |
| pig-CD31-qPCR-R1 | GGGAGCCTTCCGTTCTAGAATATC |
| pig-CD45-qPCR-F1 | CGGAGATGCAGGGTCAAACT |
| pig-CD45-qPCR-R1 | TCATCCCTTGGACCTTGTGC |
| pig-CD56-qPCR-F | GGTCAAATACCGAGCGAAGC |
| pig-CD56-qPCR-R | AGGGACTTGAGCATGACGTG |
| pig-CD29-qPCR-F | ACTTGTTGGTAAACAGCGCA |
| pig-CD29-qPCR-R | CCAGCCAATCAAAGATCCACA |
| pig-MYH3-qPCR-F | GACCGAAGACAACAGGACCC |
| pig-MYH3-qPCR-R | TGTCCTCGATCCGGTCAAAC |
| pig-MYH2-qPCR-F | GAAATCCAGCTGAACCACGC |
| pig-MYH2-qPCR-R | GGTGGATCTGGGTGTCCTTG |
| pig-MYOG-qPCR-F | TGAATGCAGTTCCCACAGCG |
| pig-MYOG-qPCR-R | GCTGTGAGCAGATGATCCCC |
| pig-ACTA1-qPCR-F | GCCGCCCTCGCCACCAGGGC |
| pig-ACTA1-qPCR-R | TACTTGAGGGTCAGGATACC |

F: Forward, R: Reverse, *: Primer is derived from "Establishment of Porcine and Human Expanded Potential Stem Cells".

**Supplementary Table.2 Antibodies used in flow cytometry or immunofluorescence**

| Antibody | Primary /Secondary | Catalogue | Brand |
|---|---|---|---|
| Anti-Fast Myosin Skeletal Heavy chain | Primary antibody | ab51263 | Abcam |
| Oct3/4 Antibody (C-10) | Primary antibody | sc-5279 | Santa Cruz Biotechnology |
| Human Nanog Antibody | Primary antibody | AF1997 | R&D systems |
| SOX2 Mouse Monoclonal Antibody | Primary antibody | EM1708-84 | HUABIO |
| PECAM-1 Alexa Fluor 488 Antibody | Primary antibody | FAB33871G | R&D systems |
| BD™ CD45 PerCP-Cy™5.5 | Primary antibody | 332784 | BD Bioscience |
| PE anti-human CD56 | Primary antibody | 985902 | BioLegend |
| Goat Anti-Mouse IgG H&L (Alexa Fluor ® 488) | Secondary antibodie | ab150113 | Abcam |
| Goat F(ab')2 Anti-Mouse IgG - Fc (PE), pre-adsorbed | Secondary antibodie | ab98742 | Abcam |
| Goat Anti-Rabbit IgG H&L (Alexa Fluor® 488) | Secondary antibodie | ab150077 | Abcam |
